# DiffuChannel: A Conditional Denoising Diffusion Model that Synthesises Single-Molecule Patch-Clamp Data with Exactly Known Idealisation (Ground Truth)

**DOI:** 10.64898/2026.10.13.757650

**Authors:** Sam T. M. Ball, Richard Barrett-Jolley

## Abstract

Development of computational models for single-channel patch-clamp analysis is hampered by limited availability of labelled (idealised) data. Since experiment alone cannot provide a genuine ground truth, two simulation methods are commonly employed; Stochastic sampling from a specified Markov model, augmented with synthetic noise allows creation of unlimited, flawlessly idealised/annotated datasets, yet its authenticity hinges entirely on the premises underlying data and noise generation. A more recent method uses GAN-based generation, this also replicates real-world noise and artefacts, but does not include a ground truth idealisation or annotation and training is notoriously unstable. Here, we introduce DiffuChannel, a conditional noise-removal diffusion model that addresses these drawbacks. Channel dynamics come directly from a user-defined discrete-time Markov chain (or a model fitted to biological seed data), thus, the state assignment for each synthetic trace is precisely determined by design. The diffusion model is then trained exclusively to capture the noise distribution of raw current conditioned on a given idealisation, thereby replicating genuine noise, filtering, and artefact patterns observed in real data without needing to hand craft noise models. We use a two-step coarse-to-fine workflow that starts with a defined slow-envelope component conditioned on longer-term dwell-time features extracted from the input idealisation. This allows a degree of baseline drift and time-dependent relaxations to be captured over timescales far longer than the network’s field of view. At generation time, we use a context-preserving inpainting method that enables creation of relatively seamless traces of any desired duration. We demonstrate that with only a limited set of labelled seed traces, synthetic records yield realistic all-points amplitude distribution, noise power spectrum, autocorrelation pattern, and slow envelope comparable to the source data. The model achieves a maximum mean discrepancy (MMD) between windowed real and generated data of 0.005 (median across the five phenotypes). This compares to a within real-split-half MMD value of 0.002 and an amplitude-matched white noise MMD value of 0.108. In this paper, we benchmark this against DeepGANnel. Since the underlying Markov model is defined rather than estimated, DiffuChannel generates infinite, perfectly labelled, phenotype-consistent training data ideal for developing and testing idealisation algorithms. We find that, in practice, this new method is far more stable than previous GAN models and runs start to end without human intervention. We provide the model open-source along with an interactive web-based training and synthesis interface/app.

## 1. Introduction

Ion channels are transmembrane proteins that control the passive movement of ions across biological membranes and contribute to the establishment of the resting membrane potential and a set of fundamental processes (Abdul Kadir et al., 2018) including the generation and propagation of action potentials (Hodgkin and Huxley, 1952). Ion channel dysfunctions give rise to a broad range of disorders called channelopathies (Sanguinetti and Spector, 1997; Jurkat-Rott et al., 2002; Buckley and Vincent, 2005; Nilius and Voets, 2013), and they are a major pharmacological target for a wide range of conditions such as pain (Staunton et al., 2013; Skerratt and West, 2015). The patch-clamp technique (Neher and Sakmann, 1976; Hamill et al., 1981; Sakmann and Neher, 1984; Neher and Sakmann, 1992) allows recording of individual channel molecular activity in real time, and single-channel recording continues to be the highest-resolution view available into the conformational dynamics of single proteins.

The main limitation in all “single channel” data analyses is arguably the initial “idealisation” process; the determination of, for every sample point in a record of potentially many millions of points, the number of channels that are open at that instant. Idealisation is usually performed by 50% threshold crossing methods or less commonly by various hidden Markov recovery approaches (Qin et al., 1996; Qin, 2004; Nicolai and Sachs, 2013; Gnanasambandam et al., 2017). To be accurate, all these methods require labour intensive expert supervision. Machine-learning approaches could potentially remove this burden, and our own Deep-Channel approach demonstrated that a hybrid convolutional–recurrent network can identify single-molecule events without human supervision (Celik et al., 2020). More recently, deep networks have been applied not to idealisation itself but to the downstream problem of recovering the kinetic scheme, by treating two-dimensional dwell-time histograms as images for classification and regression (Oikonomou et al., 2023; Oikonomou et al., 2024).

These machine learning approaches require availability of sufficient labelled/idealised training data. Generation of large datasets of real raw data is relatively easy (e.g. automated patch clamp arrays), but this includes zero ground truth labels. Supervised idealisation models need data with perfect ground truth labels at every single sample point, and hand-labelling on the scale required is just not feasible (even if the labeller could guarantee accuracy – which they cannot). Two sources of large, labelled datasets exist, and both are compromised: The first is stochastic simulation, the traditional, and probably most widely used technique. Sojourns are drawn from a hidden Markov model, typically by the Gillespie method (Gillespie, 1977) and the resulting idealisation is noised with Gaussian or similar, perhaps with a digital filter layer on top. In our Deep-Channel paper (Celik et al., 2020) we used this as the base method, but then applied authentic electrical amplifier noise by passing this signal through a genuine analogue amplifier. The labels were exact, because the idealisation was generated before the noise was applied. The weakness is that these methods do not apply authentic patch-clamp noise. The noise is hand crafted in the first case, and lacks real membrane and open channel noise in the second. Any aspect of noise, drift or other artefact the experimenter overlooks is entirely absent, and a model trained on such data risks learning the phenotype of the simulation method rather than the biological reality. Recent work has gone to considerable lengths to narrow this gap, replacing the idealised Bessel step response with an experimentally recorded one and synthesising noise from measured power spectra rather than from white noise (Oikonomou et al., 2024), work which demonstrated how much of the real signal many naive simulations miss.

The second approach is data-driven synthesis. In DeepGANnel (Ball et al., 2022) we showed that a generative adversarial network (GAN) (Goodfellow et al., 2014; Radford et al., 2015) operating on ion channel data “images” (raw current above its synchronised idealisation) can learn to emit realistically labelled records from a small annotated seed, capturing noise and artefacts that no parametric model was ever told about (Ball et al., 2022). The explicit weakness here is the opposite of the stochastic sampling/noising approach; because the GAN generates the raw trace and its label jointly, the label is also a network output. The synthetic data’s ground truth is still unknown. A GAN-generated idealisation is a plausible idealisation, but still not a true one, and a downstream idealiser trained against this could still inherit whatever systematic error the GAN made. A further, more subjective, limitation of the GAN approach is that initial training takes a long time, and the network is prone to “modal collapse” and consequently needs periodic re-starting and human monitoring (including learning rate adjustments) and inspection during training to maintain a diverse generation.

In the present paper, we attempt to resolve these issues. We use either an explicit, user-specified Markov chain to generate the idealisation or one automatically fitted to the seed data, so the ground truth is exact, controllable, and known before any network is involved, and then we apply a trained, conditional diffusion model to generate noise faithful to lab-recorded data: Open-channel and baseline noise, the amplitude and shape of the filter step response, drift, and other artefacts the seed recording contained. The generative model does not touch the idealisation/labels themselves. Idealisation is merely a *conditioning input* to the diffusion model.

For the conditional generative model, we use a denoising diffusion probabilistic model (DDPM) (Ho et al., 2020; Eschweiler et al., 2024). Diffusion models learn to gradually remove Gaussian noise, step by step. During training, they learn this by predicting the noise added to the image or signals at each stage. Diffusion methods are rapidly replacing GANs across image synthesis (Dhariwal and Nichol, 2021; Rombach et al., 2022), and showing promise in text syntheses rivalling modern large language models (LLMs) (Shen et al., 2025). Their extension to one-dimensional time series signals is now well established in domains such as audio and ECG (Chen et al., 2020; Kong et al., 2021; Rasul et al., 2021; Tashiro et al., 2021; Alcaraz and Strodthoff, 2023). We identified three advantages of DDPM over our previous GAN model (Ball et al., 2022), greater stability (e.g., no need for hand-crafted learning rates), no danger of “mode collapse” and increased training speed (whilst simulation was rapid, training itself could take days (Arjovsky et al., 2017; Ball et al., 2022).

During development of DeepGANnel, we additionally faced a challenge with any slow kinetic channel seed data; either window length needed to be increased requiring considerably more compute, or that data needed to be decimated to bring the kinetic structure into the scope of a practical window size. In the present work, we introduce a two-stage pipeline, in which a coarse model first generates a coarse or wide “envelope” from a decimated view of the idealisation, and then a fine grain model generates the detailed current conditioned jointly on the idealisation and on that coarse envelope. Combined with a context-carry inpainting scheme adapted from RePaint (Lugmayr et al., 2022), this allows generation of records of practically unlimited length without obvious window seams and with some slow drift preserved.

We believe this is the first application of a DDPM to the synthesis of patch-clamp or other single molecule data. We describe the model, the two-stage cascade, the generation procedure, and provide a set of similarity metrics. We have also made a Gradio (Abid et al., 2019) dashboard style app available that makes training and generation accessible without programming.

## 2. Methods

### 2.1. Methods summary

Our workflow can be summarised with the following steps:

1. Single-channel data consisting of a raw “noisy” current trace and its matching idealisation act as the seed for data generation.
2. This data is used to train a DPPM that maps idealisation to raw current via conditional diffusion.
3. After training the model, a discrete-time Markov chain transition matrix, defined by the user or derived from the idealisation column of the original seed data, is used to generate a synthetic idealisation of any desired length.
4. The trained DPPM model is sampled conditionally on the idealisation to yield a time-synchronized raw current.
5. The outcome is a continuous three-column dataset; Time, Channels, Noisy Current. We supply this, together with a sample of the parent data, to an analysis notebook to derive evaluation measures.

### 2.2. Data sources

Single-channel data acquisition was described previously (Ball et al., 2022), and will not be discussed further here. For this work, we have used one seed recording from each of the channel phenotypes investigated in Table 1; however, our method can be applied to any number of seed records (files) and any type of ion channel record.

**Table 1.** Data Sources. Sources of seed data used for training and evaluation. Row counts are the usable rows read from each file by the analysis notebook and are the figures quoted throughout.

| Code | Channel source | Sample size (rows) | Methods Reference |
| --- | --- | --- | --- |
| Phenotype A | Canine articular chondrocyte | 73,316 | (Lewis et al., 2013) |
| Phenotype B | WinEDR simulated | 1,000,445 | (Ball et al., 2022) |
| Phenotype C | Tracheal chondrocyte | 150,752 | (Ball et al., 2022) |
| Phenotype D | Paraventricular nucleus of the Hypothalamus | 255,840 | (Feetham et al., 2015) |
| Phenotype E | Equine articular chondrocytes | 366,588 | (Mobasheri et al., 2010), |

### 2.3. Conditional denoising diffusion theory

Note, there are two terms that conflict between the deep learning and patch-clamp literature. In computer vision deep learning, “channel” refers to the channel of an image (e.g. red, green and blue channels); whereas here a “channel” refers to the protein ion channel itself. In diffusion models, “time step” refers to each iterative noising or denoising of the data; however, in the electrophysiological field it refers to *Δt*; the elapsed time in the time series analysis. To avoid confusion, we work with a single image “channel” throughout the work, and refer to the index of the noising diffusion process as the “diffusion step”, replacing with in the conventional diffusion literature.

Let **X_0_** ∈ ℝ*^L^* be an ion channel recording of length *L* samples. Then a DDPM model defines a Markov process that gradually corrupts **X**_0_with Gaussian noise over *k* ∈ (0,N), diffusion steps:

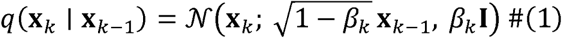

Here, *q* denotes the forward (noising) process, *N* is a multivariate Gaussian distribution and *β_k_* ∈ (0,1) the variance schedule that determines the amount of noise added at each diffusion step *k*.

It is common to reparametrize, *β_k_* into the form:

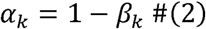

Such that:

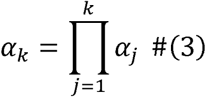

By the Markov property, this allows us to rewrite Equation 1 as:

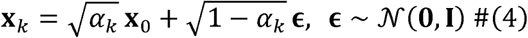

Where, **ℇ** ∈ ℝ*^L^* is a vector of independent standard Gaussian terms. This closed form allows the generation of a corruption at diffusion step without iterating through previous terms.

We then train a neural network *p_θ_* to predict the noise added by in the reverse process. We use the training loss (henceforth labelled as “eps-MSE”:

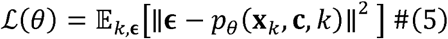

Where **c** is a conditioning signal, in this case a corresponding paired ion channel idealisation with the noisy signal **x***_k_*. The loss function is calculated over a random sample of diffusion steps *k* and noise terms **ℇ**, ensuring stability of training. This loss function is a simplified

Generation of new synthetic data works by starting with pure Gaussian noise **x***_N_* and a conditioning signal **C** to “steer” features to represent channel phenotypes:

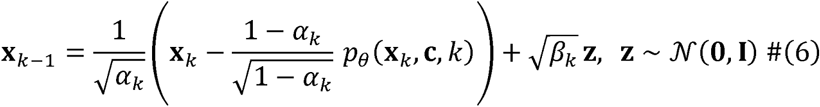

Here, **z** are randomly sampled Gaussian noise terms at each generation step; set to 0 on the **x_1_**

### 2.4. Coarse Envelope and Record Continuity

The diffusion method necessitates short, fixed window lengths due to computational feasibility of the model *p_θ_*. This excludes critical longer-term structure from the scope of the fine-grained diffusion process described above. Models trained in this way cannot simulate baseline drift, slow modulation of open-channel amplitude, or bursting structure on timescales substantially longer than its window. Furthermore, by default, when sampling from the diffusion model, outputs are independent and do not maintain continuity with previously generated samples. To mitigate this, we chose to decompose simulations across two timescales: coarse and fine, and implement continuity modelling to ensure feasible sampling across sequential generations.

#### 2.4.1. Coarse Modelling

For coarse level features (Figure 1, Figure 2), each seed record is partitioned into non-overlapping blocks of *f* samples, producing a down-sampled copy of the record. As this process reduces the sample size of windows by a factor of *f*, we repeat this process at different phase offsets to augment the training dataset size.

**Figure 1.**
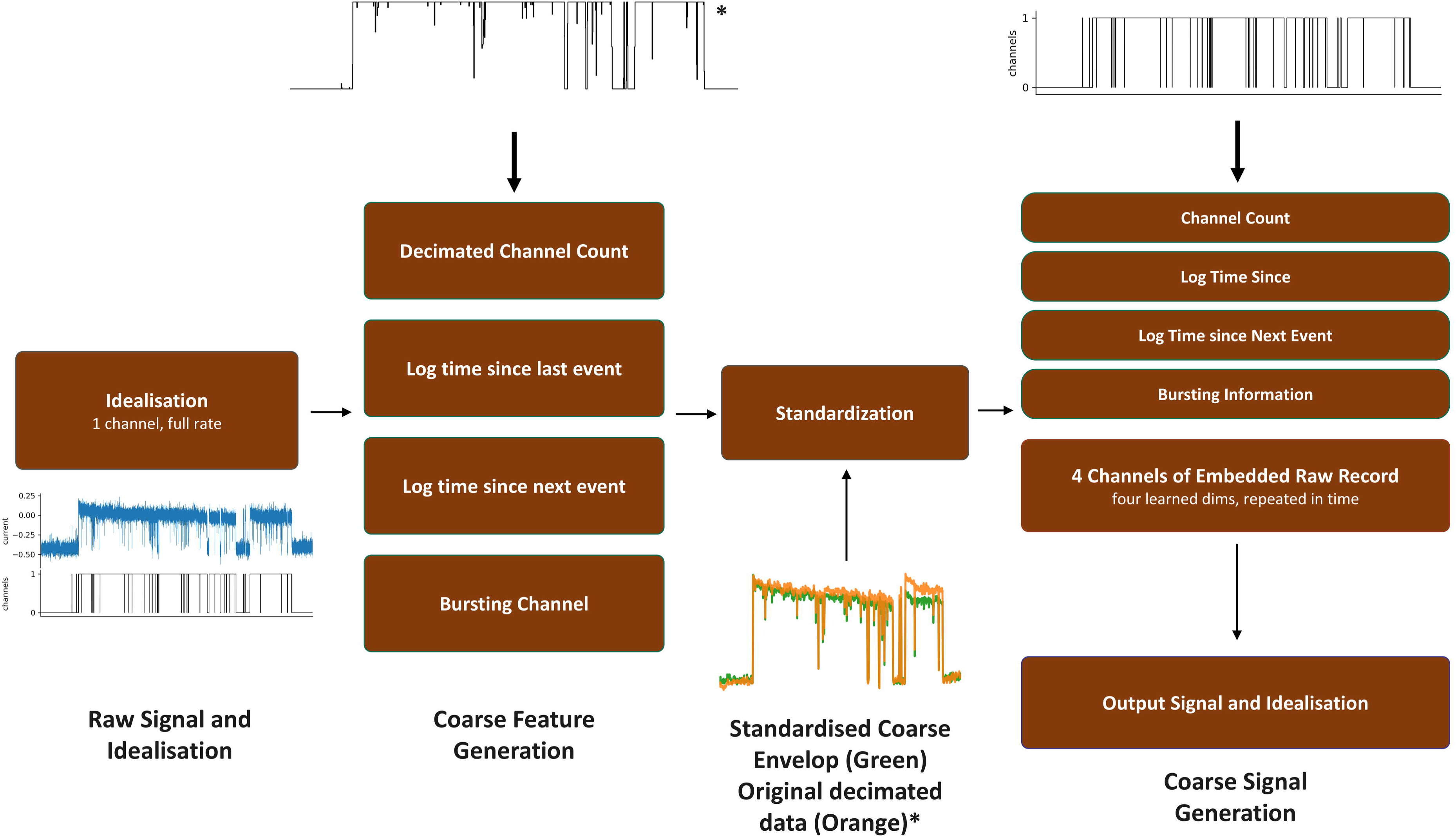
Coarse-stage input features. The coarse model receives 9 input features; the noisy data itself (which is being denoised), plus 8 conditioning features [decimated idealisation, 3 x log dwell clock features calculated *from the idealisation* (time since last transition, preceding dwell time, time since start of last burst), and a 4-dimensional feature embedding learned separately for each input seed file (this allows multi-seed input and training)]. An example noisy envelope input is shown in terms of the original seed, the decimated idealisation, then finally the slow envelope and original idealisation being fed to the fine model. *Note that the decimation procedure (block-average) creates an input idealisation in the non-integer domain even though the original was integer. **Also note the image here shows both the original (orange) alongside the actual coarse envelope (green) for illustration purposes only. The coarse envelope itself is one column of data only.

**Figure 2.**
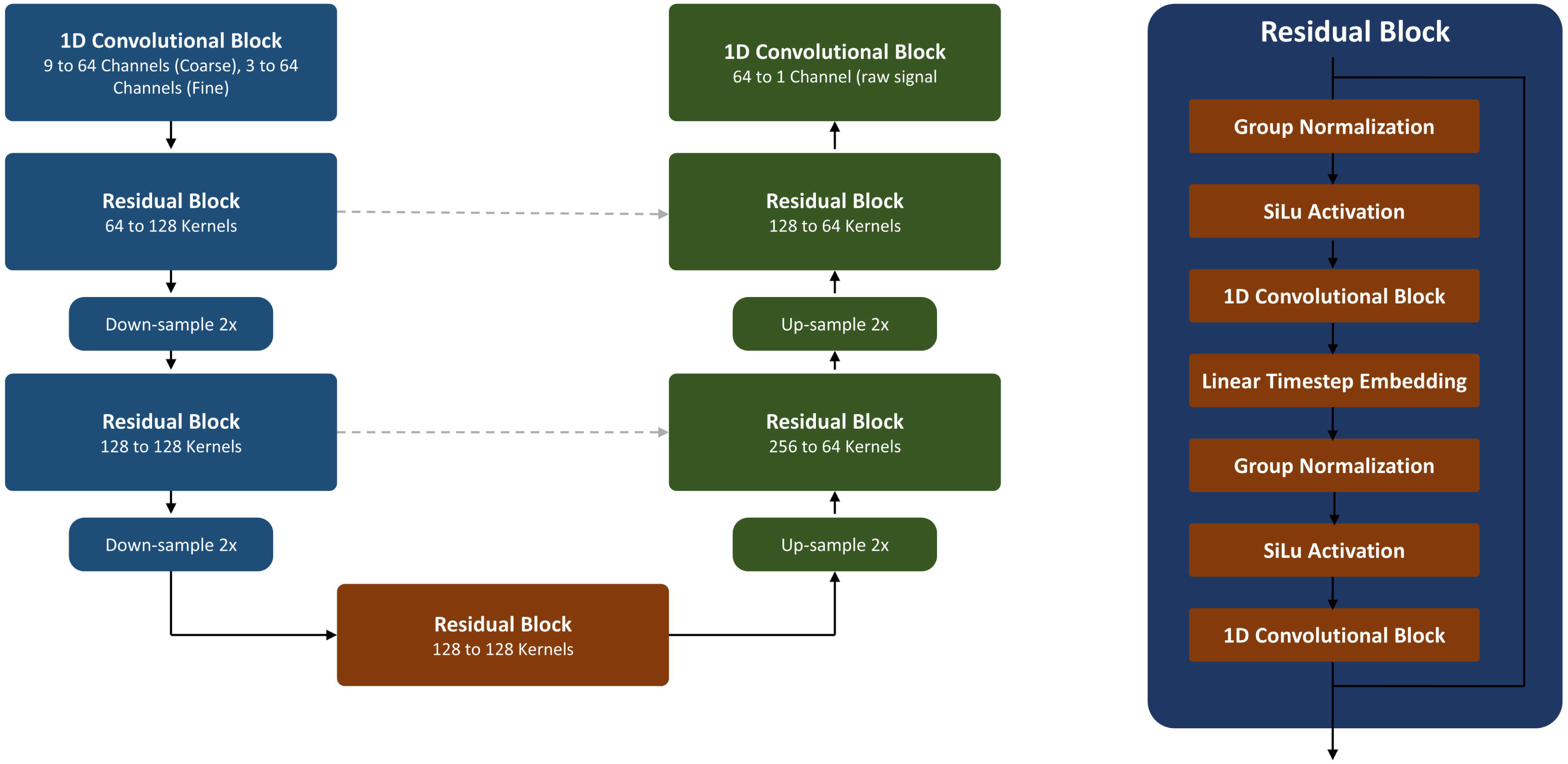
UNet1D architecture. The shared depth-2 residual U-Net used for both the coarse and fine stages; only the input convolution differs between stages (9 feature-maps for coarse, 3 for fine). Down- and up-sampling residual blocks connect through a bottleneck block and skip concatenations to a final output convolution. Parameter counts (“base_channels” = 64): 589,161 in the coarse model and 588,009 in the fine model, plus a shared 33.0k-parameter time embedding MLP feeding all five residual blocks. Note we try to avoid the usual term “channels” and prefer “feature maps” since “channels” this is confusing in an “ion channel” generation paper.

A diffusion model is trained on this view to reproduce coarse-level noise features outside of background noise. This model is conditioned on 8 channels; the first being the decimated version of the idealisation, and two channels for the log of the number of samples in the previous and current sojourn. We then use 4 further channels for an embedding of the original seed signal that is kept constant across the conditioning signal. This allows the model to associate specific features with each seed record across coarse windows. For training, we include Dropout with probability factor of 0.1, and for generation this embedding is supplied to allow for the characteristic noise for the entire record to be reproduced. The final channel is reserved for the burst events; we log the time elapsed since the start of the current burst (if any), ignoring any sojourns shorter than 1000 samples to simulate characteristic “bursting” behaviour of ion channels in single channel recordings.

These features are all present in the base idealisation, so do not lead to data “leakage”. They were engineered as examples of physiologically and structurally important features that would result in higher quality outputs.

After generation, this single channel envelope is standardised and expanded back to full sample rate by repetition and used as a second conditioning channel for the fine model below.

#### 2.4.2. Fine Model

The fine model (Figure 2) now has access to a broader context via its conditioning, and is trained with this coarse feature channel as conditioning. The model is trained as above; predicting the noise of samples of channels, but with the additional information of long-term features in the underlying recording (Figure 3).

**Figure 3.**
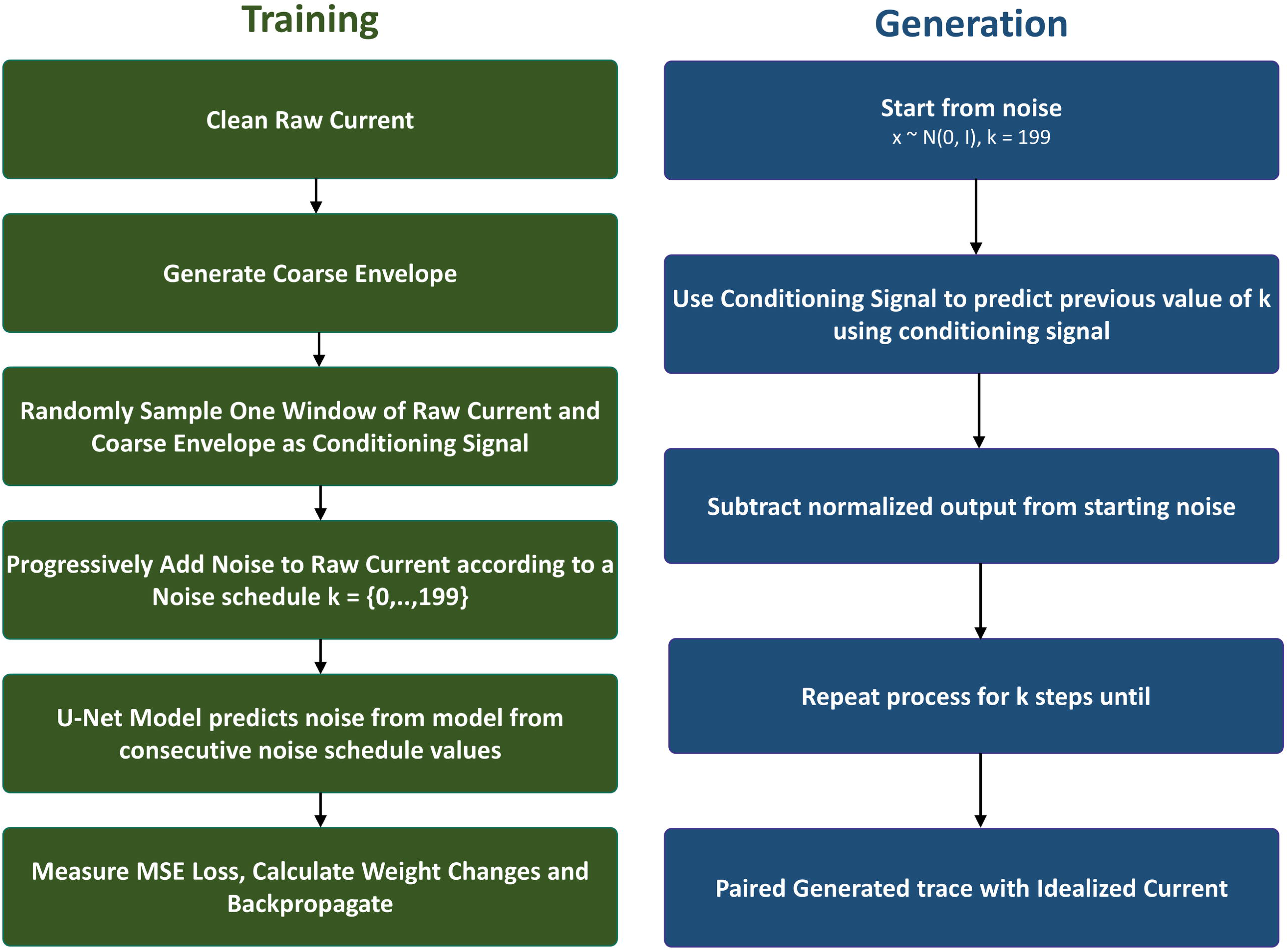
The shared diffusion process. Identical forward and reverse diffusion mathematics are used for both stages (betas linearly spaced from 1e-4 to 0.02 over 200 steps); only the conditioning channels differ between them. Training: a diffusion step k is drawn uniformly from 0-199, the corresponding noised window x_k is formed, and the network is trained by MSE loss to predict the noise that was added. Generation: starting from Gaussian noise, the network’s predicted noise is subtracted at each of 200 reverse steps (one full U-Net pass per step, per window), with the conditioning channels concatenated to x_k throughout but never themselves noised. The coarse model’s output is the generated envelope; the fine model’s output is the generated current.

#### 2.4.3. Continuity Assurance Across Samples

Long sequences quickly result in infeasible model sizes for training and generation, as model size scales non-linearly with window length. Traditionally, the approach for channel simulation would be to simulate non-overlapping sections of the record and join them together at the edges; however, this results in extreme discontinuities in the record where the underlying idealisations do not match.

To mitigate this, we generate records in overlapping blocks and blend blocks by overwriting portions of subsequent regions in the diffusion process known as the RePaint context-carry scheme (Lugmayr et al., 2022); applied here on the time axis. At every diffusion generation step *k*, we overwrite the first 512 samples of the block with the previously generated block’s final 512 samples. This ensures continuity between blocks, and allows the model to use this previous information as a source for diffusion in future blocks.

### 2.5. Fitting and Simulation of the Conditioning Idealisation

Idealisations are generated using a discrete-time hidden Markov chain with a per-sample transition probability matrix connecting many “hidden” states (thought to correlate with physical configurations of the protein itself) with “visible” states (opening and closing of the channel itself) (Colquhoun and Hawkes, 1981; Fredkin and Rice, 1986). *P* governs the kinetic behaviour of the ion channel and has been shown to have pharmacological implications; it is likely that pharmacological agents change the nature of this underlying matrix to effect ion channel behaviour. The correct choice of *P* is therefore important, and we do so by fitting to the idealisation of a seed record before generation so that simulated data matches both the noise and kinetic structure of the underlying data. We fit *P* using the Baum-Welch algorithm (Baum et al., 1970; Rabiner, 1989), which is simplified in this case as each hidden state corresponds with exactly one of the visible states.

Presence of more than one channel in a simulated patch creates additional complications (*n=*number of channels in the patch is a settable parameter in our generation phase, and should ideally match whatever was in the original seed record). To address this, we assume firstly that channels are kinetically identical (same HMM matrix and channel amplitude), and we then model the whole patch as every possible combination of individual channel states. To fit this model robustly, we start the fitting from three different initial guesses and add a small statistical correction so that rarely-seen transitions are not entirely eliminated. The run with the highest log-likelihood is retained. The Bayesian information criterion (BIC) (Schwarz, 1978), which penalises unnecessary complexity, is reported for every fit so that alternative topologies can be compared; all results here use the single four-state topology shown in Table 2.

**Table 2.** Transition matrices. Transition probabilities extracted from fits to each seed record and then used for data generation (conditioning idealisation. “P12” is the probability of moving from state 1 to state 2 etc. Note that impossible transitions were constrained with zero and are not shown in the table. For example, there is no direct connection from state 1 to state 4. States 1 and 2 are “closed” (non-conducting states) whereas 3 and 4 are “open”.

| Record | p11 | p12 | p21 | p22 | p23 | p32 | p33 | p34 | p43 | p44 |
| --- | --- | --- | --- | --- | --- | --- | --- | --- | --- | --- |
| Topology (start) | 0.970 | 0.030 | 0.020 | 0.900 | 0.080 | 0.100 | 0.850 | 0.050 | 0.200 | 0.800 |
| Phenotype A | 0.997 | 0.003 | 0.006 | 0.906 | 0.089 | 0.068 | 0.881 | 0.051 | 0.020 | 0.980 |
| Phenotype B | 0.988 | 0.012 | 0.087 | 0.908 | 0.005 | 0.010 | 0.899 | 0.091 | 0.175 | 0.825 |
| Phenotype C | 0.999 | 0.001 | 0.014 | 0.928 | 0.058 | 0.023 | 0.895 | 0.083 | 0.004 | 0.996 |
| Phenotype D | 0.385 | 0.615 | 0.002 | 0.737 | 0.261 | 0.005 | 0.913 | 0.082 | 0.141 | 0.859 |
| Phenotype E | 0.983 | 0.018 | 0.011 | 0.877 | 0.111 | 0.012 | 0.983 | 0.005 | 0.035 | 0.965 |

### 2.6. Neural Network architecture

For our choice of denoising model,*P_θ_*, we use a one-dimensional U-Net model (Ronneberger et al., 2015) for both fine and coarse modelling (Figure 2). The total parameter count for the combined model was 1,177,170 parameters with 588,009 in the fine model and 589,161 parameters in the coarse model. For training, the diffusion step index is embedded with sinusoidal encoding (Vaswani et al., 2017) and passed through a two-layer linear model with SiLU activation (Elfwing et al., 2018). The resulting embedding is projected and added to the feature map inside every residual block. Each residual block applies group normalisation (Wu and He, 2018) with 8 groups, SiLU, and a 3-tap convolution, twice, with an identity residual connection. The encoder path is a residual block at 64 channels followed by a strided convolution down sampling by 2, then a residual block at 2×64 channels followed by a second down sampling; a bottleneck residual block; then a symmetric decoder path of transposed convolutions with skip concatenation from the matching encoder stage. A final GroupNorm–SiLU–convolution emits a single-channel noise prediction. Window lengths are constrained to multiples of four by the two down sampling stages.

### 2.7. Model Hyperparameters

We use a linear variance schedule,*β_k_* increasing linearly from *β*_1_10^-4^ to β*_k_*=2×10^-2^ (Ho et al., 2020), with *K*= 200 diffusion steps. We used AdamW optimisation (Loshchilov and Hutter, 2019) with learning rate and weight decay both set to 0.0001 and a batch size of 32. Training ran for 20 epochs at the fine stage, but 80 for the coarse envelope to compensate for the 64-fold reduction in available windows after decimation.

### 2.8. Evaluation metrics

We are not aware of a previous attempt to simulate patch clamp data via a diffusion model and therefore we compare real, seed data records to diffusion generated data alongside real to our previous GAN generated data comparisons. Traces are cropped to the minimum length for a given ion channel phenotype in order to avoid one obvious source of bias. Benchmarking metrics were performed with R. We measure several diffusion and electrophysiological metrics/tests for model performance including:

i) Maximum mean discrepancy (MMD) (Gretton et al., 2012), a statistical measure of difference between two probability distributions by considering *X,Y* by considering *N_X_* samples of **x**_i̴̴_ ∼ *X:*

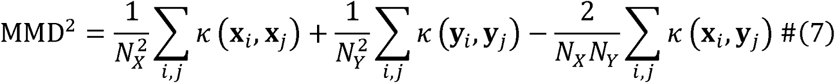 Where *k is* a Gaussian Kernel:

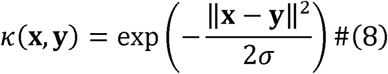 Where *N* is the sample size and σ is the kernel bandwidth. In theory, the MMD of samples taken from identical distributions tends to 0 as the size of each sample increases.
ii) In electrophysiology, it is more common to compare all-points amplitude histograms in single channel experiments for changes to channel function. We also therefore additionally test for similarity of kernel density estimates (KDE) of the full sample distributions using a two-sample Kolmogorov-Smirnov test.
iii) Another physiologically important measure is the ability to reproduce data that replicates the dwell time histogram (Sigworth and Sine, 1987) of the seed data. We therefore run another Kolmogorov-Smirnov test on the KDE distributions of these histograms to test model accuracy here.
iv) Perhaps the most directly sensitive measure for physiology is the reproduction of authentic noise from seed data. Oikonomou and colleagues have shown this to be badly misrepresented by conventional simulation (Oikonomou et al., 2024) and previous methods. A Welch transformation (Welch, 1967) of the data allows for an additional analysis of the noise power spectral density distribution, for which we run an additional Kolmogorov-Smirnov test to check similar distribution of simulated output to seed data.
v) Finally, as in DeepGANnel, we use manifold projections (van der Maaten and Hinton, 2008; McInnes et al., 2018) of 512 sample windows to check the distribution of records in low dimensional space using permutation tests (ClusterSignificance (Serviss et al., 2017) in R).

### 2.9. Software and Hardware

DiffuChannel is written in Python with two modules; the main model in one script and the Markovian fitting algorithms in a second script. We deploy a browser-based dashboard interface using the Gradio package (Abid et al., 2019). This can be run without any scripting/programming. The app displays loss curves and example online during training along with amplitude histograms of seed and generated etc. Generated records are saved in parquet pandas data-frame format with Time, Channels and Noisy Current columns. Training was run on a single CUDA GPU (PyTorch 2.6.0+CUDA 31.1, Nvidia RTX Titan).

All figures and metrics reported here can be produced by running the associated R-notebook. GAI (Claude) was used to assist with coding, debugging and editing of the manuscript itself.

## 3. Results

### 3.1. Training Progress

Seed records (Table 1) were idealised, standardised and windowed, yielding 115,395 training windows in total: 4,575, 62,520, 9,414, 15,982 and 22,904 for phenotypes A to E respectively. The conditional diffusion model was trained for 20 fine and 80 coarse envelope epochs. Training and generation times were broadly similar across phenotypes A to E, with the total diffusion training time taking around 2.2 hours. Markov fitting added 22 to 104 seconds to the training processes depending on the complexity of the model. Figure 4 show the evolution of the eps-MSE objective during training, for both the coarse and fine stages of the model, across different phenotypes.

**Figure 4.**
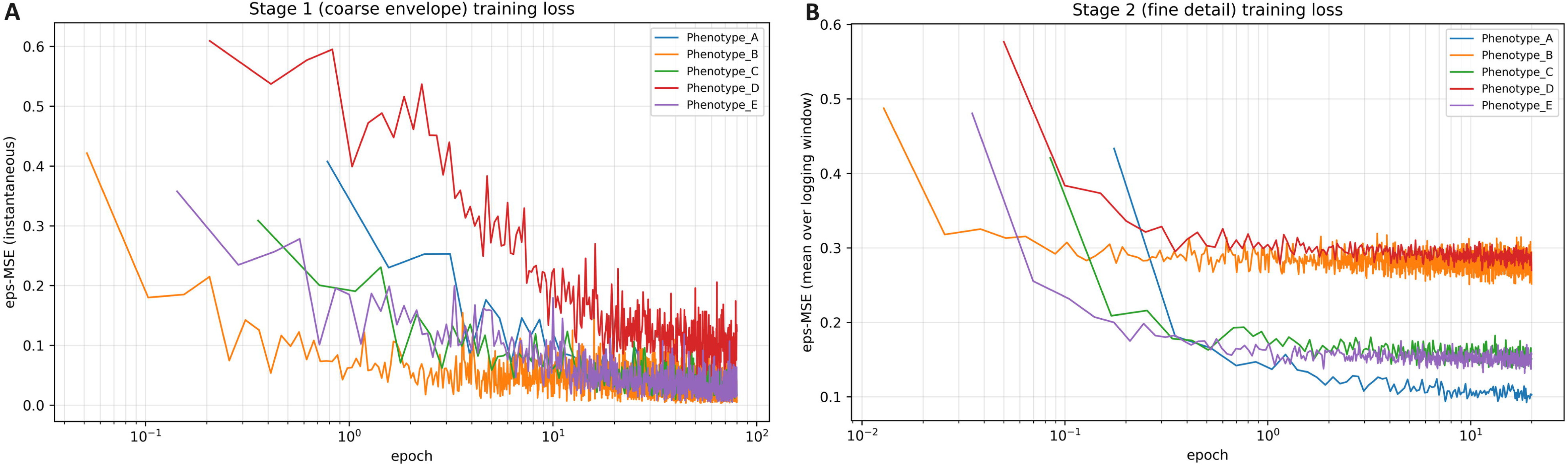
ε-MSE Training losses. ε-MSE training loss against epoch for **(A)** the coarse and **(B)** fine stages, log abscissa, one line per phenotype. We use ε-MSE training loss (ie., difference of Gaussian noise: *mean((ε^* − ε))) to distinguish from an MSE of ((original - generated)) since the intention is to reproduce the noise structure not an identical record. We ran all these experiments with 20×4 epochs course. Steps per epoch scaled with record length. This was an arbitrary choice and there was no early stopping applied. Clearly Stage 1 coarse training was still improving at 20×4 epochs for Phenotype D, for example.

### 3.2. Qualitative Analysis

Figure 5A shows outputs of the diffusion model alongside their paired seed and GAN generated signal, Figure 5B shows the underlying idealisation (conditioning or generated) from the same records as that in Figure 5A. Subjectively we find that the generated records show the characteristic features of the phenotype, including noise texture, transition envelopes from recording filters, and increased open-channel noise (Sigworth, 1985).

**Figure 5.**
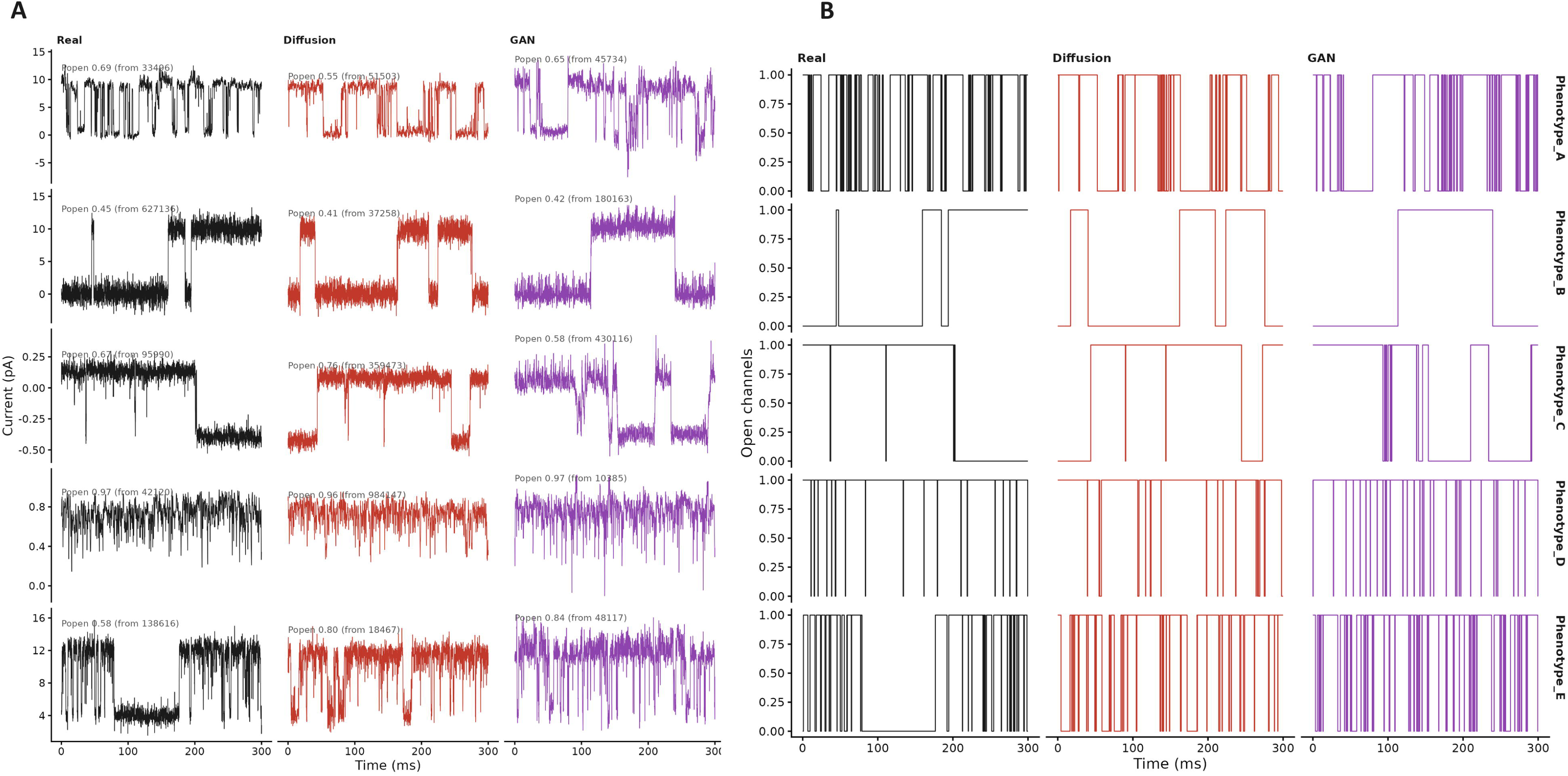
Representative data. Example 300 ms traces: real, diffusion and DeepGANnel, one row per phenotype, with the open probability of the window shown. **(A)** is the Noisey Current, and **(B)** is the idealisation either from the seed data or generated records.

### 3.3. Distributional Analysis

For a more quantitative approach, we record the maximum mean discrepancy of the records and a Kolmogorov-Smirnov test for the all-points amplitude histogram.

Figure 6 shows the all-points amplitude histograms for real and generated data (as kernel density plots - to allow fully automated fitting). We find that single level noise deviations look subjectively similar to their corresponding seed data. Tables 3 and 4 quantify the difference/similarity. Synthetic data from both synthetic arms look subjectively realistic, and its amplitude distribution was statistically equivalent to that of the seed record within the real record’s own split-half margin (Table 3): equivalence was confirmed in all four out of five phenotypes for DiffuChannel (for the fifth phenotype D, p-value was a little over 0.05, (0.052)) and in four of five for DeepGANnel. Mean equivalence-test p-values for DiffuChannel to Real and GAN to real were 0.044 (range 0.030 to 0.052) and 0.042 (0.024 to 0.076) respectively. The maximum mean discrepancy (MMD) was calculated (equation 7) for windowed samples between the real vs GAN generated and Real v Diffusion generated data. Table 5 gives the MMD of each synthetic arm against its seed record for every phenotype; the median across phenotypes was 0.005 for DiffuChannel and 0.008 for DeepGANnel, against 0.002 for the within-real split-half floor and 0.108 for amplitude-matched Gaussian white noise.

**Figure 6.**
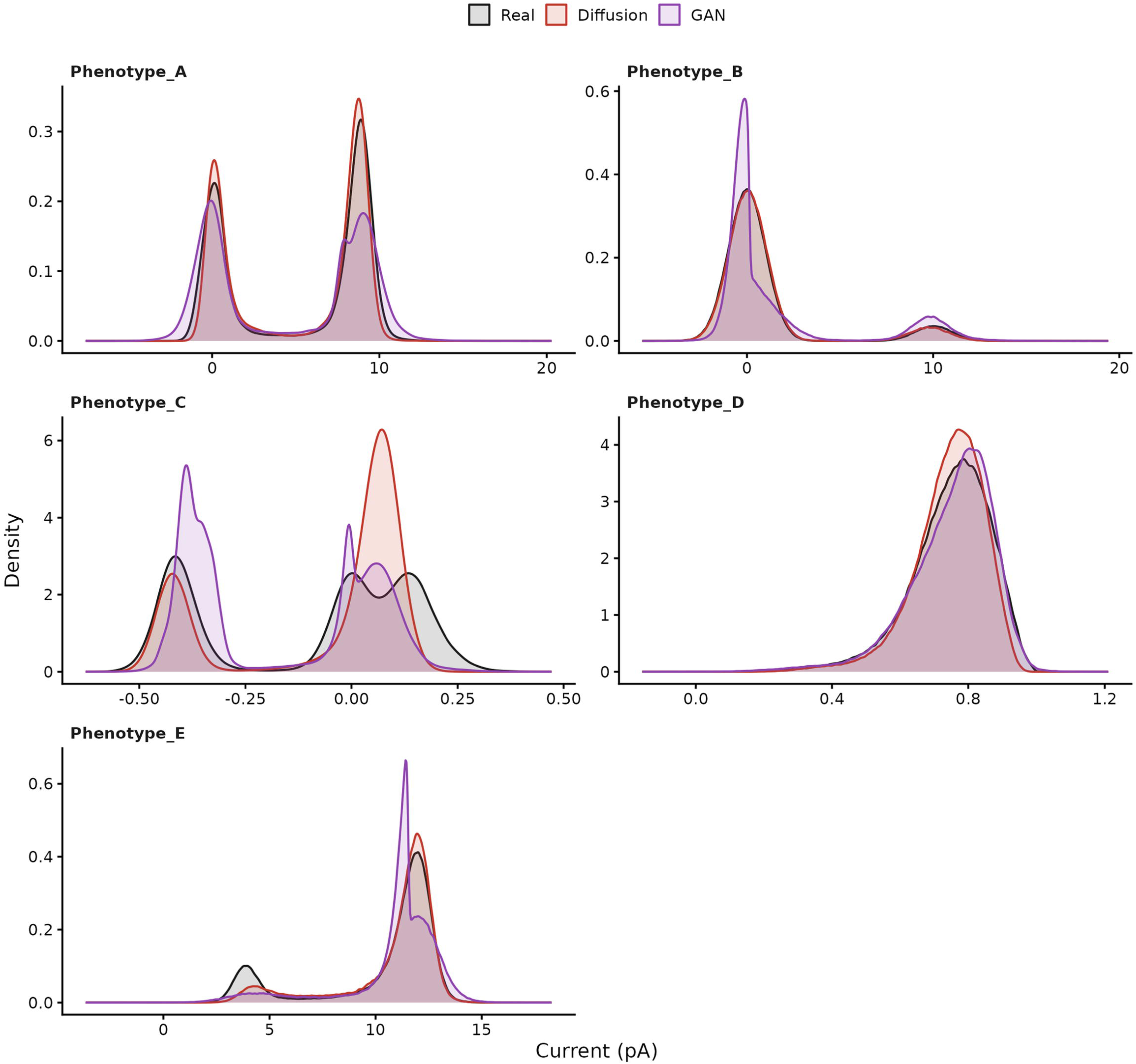
All-points amplitudes. Three overlapping kernel density plots for each phenotype used. One plot for each arm; real record (black), the diffusion-generated record (red) and the DeepGANnel-generated record (purple). Note that rather than the conventional binned and properly fitted amplitude histograms, these are shown simply as smoothed density estimates (kernel density, bandwidth on a per record basis, between 0.003 and 0.25 pA (1–6% of each record’s standard deviation), see R benchmarking script. This allowed a fully automated benchmarking process.

**Table 3.** Amplitude equivalence. Nonparametric KS-distance (windowed). One-sided equivalence test (Wellek, 2010) on the Kolmogorov-Smirnov (KS) distance between windows of the current trace (1024 samples). Δ is the equivalence margin, set by the variance within the real record alone as the 95th percentile of the KS distance between windows drawn from real’s own first and second halves. *p* is the Wellek *p-value* for equivalence (lower value is less likely to be equivalent through chance alone). *Upper bound* is the 95th percentile of the KS-distance distribution between real and arm windows. *Verdict* is ‘Equivalent’ if Upper bound < Δ, or ‘Not confirmed equivalent’ otherwise. *Winner* is simply the arm confirmed equivalent when only one of the two is; ‘Tie’ when both or neither are.

| Phenotype | Arm | Window length (n samples) | $\Delta$ | Upper bound | $p$ | Verdict | Winner |
| --- | --- | --- | --- | --- | --- | --- | --- |
| A | Diffusion | 1024 | 0.87 | 0.84 | 0.044 | Equivalent |  |
| A | GAN | 1024 | 0.87 | 0.87 | 0.046 | Equivalent | **Tie** |
| B | Diffusion | 1024 | 0.50 | 0.48 | 0.048 | Equivalent |  |
| B | GAN | 1024 | 0.50 | 0.55 | 0.076 | Not confirmed equivalent | **Diffusion** |
| C | Diffusion | 1024 | 1.00 | 1.00 | 0.046 | Equivalent |  |
| C | GAN | 1024 | 1.00 | 1.00 | 0.042 | Equivalent | **Tie** |
| D | Diffusion | 1024 | 0.26 | 0.26 | 0.052 | Equivalent |  |
| D | GAN | 1024 | 0.26 | 0.24 | 0.024 | Equivalent | **Tie** |
| E | Diffusion | 1024 | 0.92 | 0.74 | 0.030 | Equivalent |  |
| E | GAN | 1024 | 0.92 | 0.75 | 0.024 | Equivalent | **Tie** |

**Table 4.** Noise-SD distribution equivalence. One-sided equivalence test (Wellek, 2010) on the distribution of current SD measured per qualifying open/closed event (≥30 consecutive samples in one state. *n* (real/arm) is the number of qualifying events contributing to real windows and that of the arm. Δ is the 95th percentile of the KS distance between real’s own first-half and second-half events (fixed from real alone, as Table 3). *Upper bound* is the 95th percentile of the real-vs-arm KS-distance distribution. *Verdict* is ‘Equivalent’ if Upper bound < Δ. *Winner* is the arm confirmed equivalent when only one of the two is; ‘Tie’ when either both or neither are.

| Phenotype | Arm | State | n<br>(real/arm) | $\Delta$ | Upper<br>bound | $p$ | Verdict | Winner |
| --- | --- | --- | --- | --- | --- | --- | --- | --- |
| A | GAN | Open | 330/382 | 0.20 | 0.52 | 1.000 | Not confirmed equivalent |  |
| A | Diffusion | Open | 330/428 | 0.20 | 0.22 | 0.134 | Not confirmed equivalent | **Tie** |
| A | GAN | Closed | 141/123 | 0.32 | 0.31 | 0.030 | Equivalent |  |
| A | Diffusion | Closed | 141/134 | 0.32 | 0.23 | 0.002 | Equivalent | **Tie** |
| B | GAN | Open | 399/281 | 0.34 | 0.27 | 0.002 | Equivalent |  |
| B | Diffusion | Open | 399/228 | 0.34 | 0.36 | 0.102 | Not confirmed equivalent | **GAN** |
| B | GAN | Closed | 460/299 | 0.27 | 0.16 | 0.002 | Equivalent |  |
| B | Diffusion | Closed | 460/271 | 0.27 | 0.19 | 0.002 | Equivalent | **Tie** |
| C | GAN | Open | 82/1279 | 0.49 | 0.31 | 0.002 | Equivalent |  |
| C | Diffusion | Open | 82/542 | 0.49 | 0.35 | 0.002 | Equivalent | **Tie** |
| C | GAN | Closed | 57/376 | 0.51 | 0.42 | 0.004 | Equivalent |  |
| C | Diffusion | Closed | 57/212 | 0.51 | 0.40 | 0.004 | Equivalent | **Tie** |
| D | GAN | Open | 762/5889 | 0.12 | 0.27 | 1.000 | Not confirmed equivalent |  |
| D | Diffusion | Open | 762/2864 | 0.12 | 0.46 | 1.000 | Not confirmed equivalent | **Tie** |
| D | GAN | Closed | N/A | N/A | N/A | N/A | Skipped (too few qualifying real events for a margin) |  |
| D | Diffusion | Closed | N/A | N/A | N/A | N/A | Skipped (too few qualifying real events for a margin) | **N/A** |
| E | GAN | Open | 2143/1858 | 0.14 | 0.12 | 0.002 | Equivalent |  |
| E | Diffusion | Open | 2143/1874 | 0.14 | 0.07 | 0.002 | Equivalent | **Tie** |
| E | GAN | Closed | 344/204 | 0.26 | 0.33 | 0.497 | Not confirmed equivalent |  |
| E | Diffusion | Closed | 344/258 | 0.26 | 0.48 | 1.000 | Not confirmed equivalent | **Tie** |

**Table 5.** Similarity metrics. Further, generic similarity metrics, computed identically for both synthetic arms (Diffusion and GAN) against the real seed record of each phenotype (A–E) on the majority class of that phenotype. Lower values are better unless stated. *ACF RMS*: root-mean-square (RMS) difference between the autocorrelation functions (ACF) of real and synthetic noise at lags of 0–200 samples, measured on single-level segments (closed level for phenotypes A–C, open level for D and E; as Figure 7B). *Log10 PSD RMS*: RMS difference between the log₁₀ Welch power spectral densities (PSD) of the same segments, over a common frequency grid up to the 1 kHz filter corner (as Figure 7A). *KS amplitude*: two-sample Kolmogorov–Smirnov (KS) statistic *D*, the largest vertical distance between the cumulative all-points amplitude distributions of real and synthetic current (Figure 6). *MMD vs real:* maximum mean discrepancy (MMD) between 400 randomly drawn 128-sample windows of real and of synthetic current (whose width was set to the median distance between windows). Zero means indistinguishable; small negative values can occur by chance. *MMD permutation p*: permutation-test p-value for that MMD, from 200 random relabellings of the pooled windows as real or synthetic. A large p means the real–synthetic difference is not statistically greater than simple relabelling produces. Here, higher implies the data is more similar; the smallest possible value is 1/201 ≈ 0.005. *MMD envelope:* MMD as above, computed only on the slow envelope (decimated with block means of 64 samples) in windows of 64 blocks. *Envelope SD and Noise SD (pA):* standard deviation of the slow envelope, and of the current at the single analysed conductance level. Open probability and Mean open/closed dwell (ms): from each arm’s Channels column. For these five, the Real (reference) column gives the real seed record’s own value, and thus the arm closer to this value is better.

| Metric | Phenotype | Diffusion | GAN | Real (reference) |
| --- | --- | --- | --- | --- |
| ACF RMS | A | 0.074 | 0.105 | - |
| ACF RMS | B | 0.003 | 0.016 | - |
| ACF RMS | C | 0.027 | 0.164 | - |
| ACF RMS | D | 0.035 | 0.040 | - |
| ACF RMS | E | 0.012 | 0.020 | - |
| Envelope SD (pA) | A | 3.433 | 3.998 | 3.639 |
| Envelope SD (pA) | B | 2.549 | 3.424 | 2.685 |
| Envelope SD (pA) | C | 0.204 | 0.205 | 0.239 |
| Envelope SD (pA) | D | 0.042 | 0.048 | 0.052 |
| Envelope SD (pA) | E | 1.625 | 1.465 | 2.414 |
| KS amplitude | A | 0.061 | 0.099 | - |
| KS amplitude | B | 0.015 | 0.097 | - |
| KS amplitude | C | 0.161 | 0.195 | - |
| KS amplitude | D | 0.058 | 0.035 | - |
| KS amplitude | E | 0.082 | 0.117 | - |
| MMD envelope | A | 0.010 | 0.027 | - |
| MMD envelope | B | 0.000 | 0.037 | - |
| MMD envelope | C | 0.025 | 0.109 | - |
| MMD envelope | D | 0.022 | 0.006 | - |
| MMD envelope | E | 0.014 | 0.026 | - |
| MMD permutation p | A | 0.677 | 0.005 | - |
| MMD permutation p | B | 0.109 | 0.005 | - |
| MMD permutation p | C | 0.005 | 0.005 | - |
| MMD permutation p | D | 0.005 | 0.005 | - |
| MMD permutation p | E | 0.005 | 0.005 | - |
| MMD vs real | A | -0.001 | 0.010 | - |
| MMD vs real | B | 0.000 | 0.003 | - |
| MMD vs real | C | 0.042 | 0.046 | - |
| MMD vs real | D | 0.007 | 0.003 | - |
| MMD vs real | E | 0.005 | 0.008 | - |
| Mean closed dwell (ms) | A | 3.691 | 4.485 | 3.401 |
| Mean closed dwell (ms) | B | 157.848 | 123.969 | 195.175 |
| Mean closed dwell (ms) | C | 28.687 | 14.068 | 37.275 |
| Mean closed dwell (ms) | D | 0.388 | 0.220 | 0.380 |
| Mean closed dwell (ms) | E | 0.739 | 0.534 | 1.195 |
| Mean open dwell (ms) | A | 5.163 | 4.976 | 4.751 |
| Mean open dwell (ms) | B | 15.669 | 22.567 | 19.094 |
| Mean open dwell (ms) | C | 85.733 | 13.272 | 70.912 |
| Mean open dwell (ms) | D | 31.158 | 14.802 | 30.897 |
| Mean open dwell (ms) | E | 4.651 | 4.160 | 5.138 |
| Noise SD (pA) | A | 1.378 | 1.551 | 1.438 |
| Noise SD (pA) | B | 1.214 | 1.246 | 1.174 |
| Noise SD (pA) | C | 0.040 | 0.056 | 0.048 |
| Noise SD (pA) | D | 0.096 | 0.114 | 0.109 |
| Noise SD (pA) | E | 0.999 | 1.120 | 1.022 |
| Open probability | A | 0.583 | 0.526 | 0.583 |
| Open probability | B | 0.090 | 0.154 | 0.089 |
| Open probability | C | 0.749 | 0.485 | 0.654 |
| Open probability | D | 0.988 | 0.985 | 0.988 |
| Open probability | E | 0.863 | 0.886 | 0.811 |
| log10 PSD RMS | A | 0.544 | 0.554 | - |
| log10 PSD RMS | B | 0.151 | 0.513 | - |
| log10 PSD RMS | C | 0.308 | 0.892 | - |
| log10 PSD RMS | D | 0.195 | 0.155 | - |
| log10 PSD RMS | E | 0.081 | 0.075 | - |

### 3.4. Noise and Filter Response Analysis

Power Spectrum Density (PSD) analysis via Welch’s estimates tested for the reproduction of convincing noise in generated data compared to the seed data it was produced from, an area that has been consistently challenging previously as shown by Oikonomou and colleagues (Oikonomou et al., 2024). Figure 7A shows the Welch PSD estimates for the single-state segments of real and generated records. We find that the two spectra match closely across all frequency bands, instead of producing a profile consistent with “plain” white noise that is often used in ion channel simulation pipelines. The RMSE between the log spectra had a mean of 0.26 units from the diffusion arm compared to 0.44 for DeepGANnel, with the diffusion arm closer in three of the five phenotypes (Table 5). Above 1kHz the DeepGANnel spectra contain excessive high-frequency noise that the real record does not, that our new method emulates well. Figure 7B shows the similarities between local noise structure across the 3 data types (real, GAN and diffusion) for each of the 5 phenotypes. This is illustrated with autocorrelation analysis; correlating how similar data is to itself after a short delay. In all 5 phenotypes, the Diffusion model follows the real data more closely that DeepGANnel.

**Figure 7.**
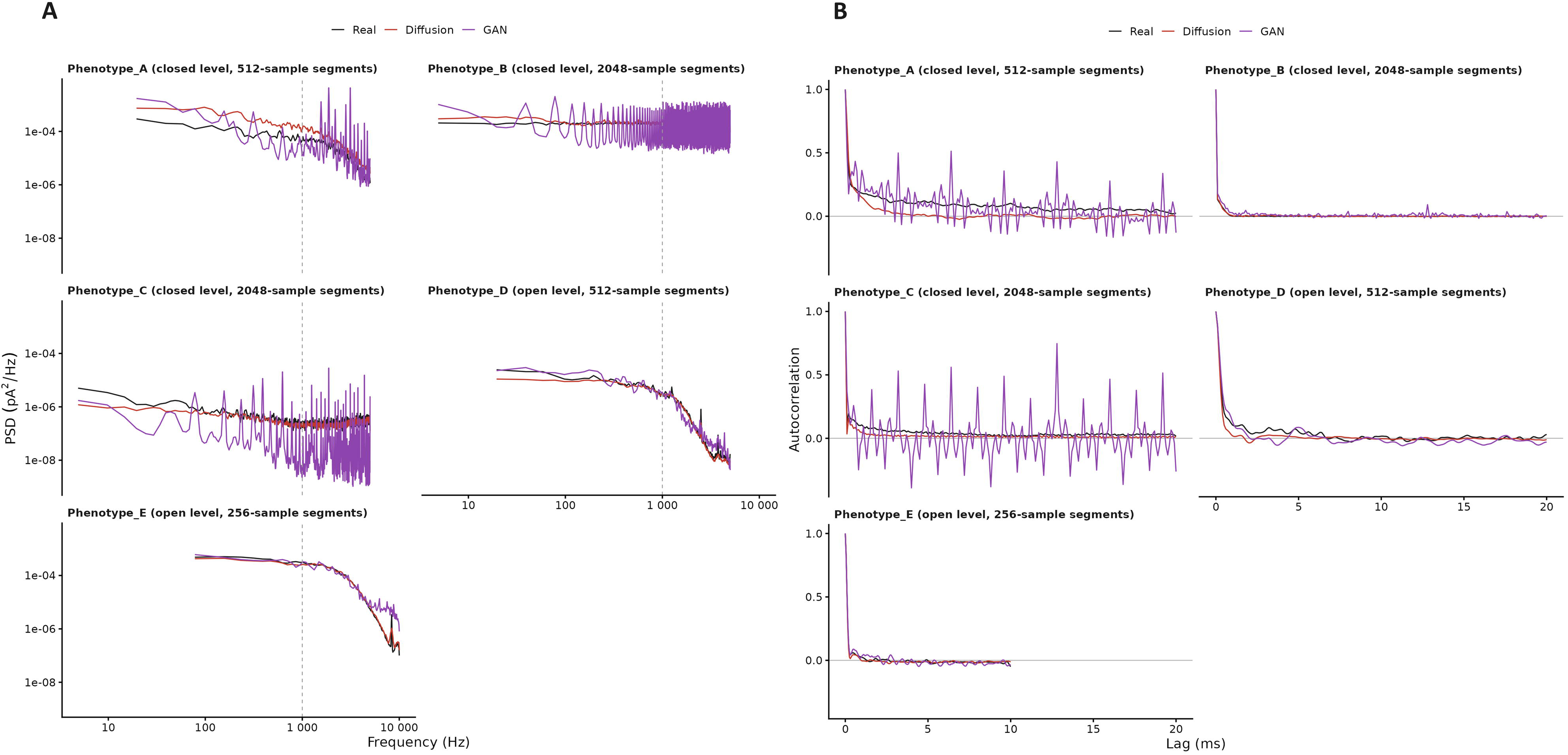
Noise structure, of the Noisy Current signal comparing Real, GAN and Diffusion across all 5 phenotypes. A single conductance level is analysed per phenotype, chosen as the level at which all three arms supply the longest uninterrupted (transition-free) segments, since a spectrum can only be estimated within a single level. Closures in phenotypes D and E are too brief for this (the longest closure in at least one arm is 6 and 149 samples respectively, below the 256-sample minimum segment). **(A)** Welch power spectral density of single-level segments for all three arms, log-log, one panel per phenotype. The conductance level and segment length used are stated in each panel: the closed level for phenotypes A to C and the open level for D and E. The dashed line marks the 1 kHz filter corner. **(B)** Autocorrelation analyse of the same segments as part A. Autocorrelation measures how similar the data points are to datapoints from the same sample after a given time lag. All curves begin at 1 on the y-axis with zero lag (perfect correlation) and approach zero (no correlation) as the lag increases. Across all five phenotypes, the diffusion model curve (red) more closely follows the real-data curves (black) than those of DeepGANnel (purple), indicating that it better reproduces the short-term noise structure.

### 3.5. Coarse envelope

Figure 8A shows decimated copies (factor 64) of both the real record and coarse-stage generated records from our diffusion network alongside equally decimated GAN data, and Figure 8B shows the distribution of envelope (amplitude histogram - kernel density plots) for the real, GAN and Diffusion models over the same periods. The standard deviation of the envelope in the simulated records compared to their seed is shown in Table 5, with the diffusion records showing a lower MMD compared to the GAN based method when compared to the seed distribution. This suggest that the diffusion method not only more accurately reproduces noise consistent with the underlying seed record, but also better captures longer-term trends in the recording such as single channel current decay (Phenotype C), a common real world phenomenon in single channel recording (Fenwick et al., 1982).

**Figure 8.**
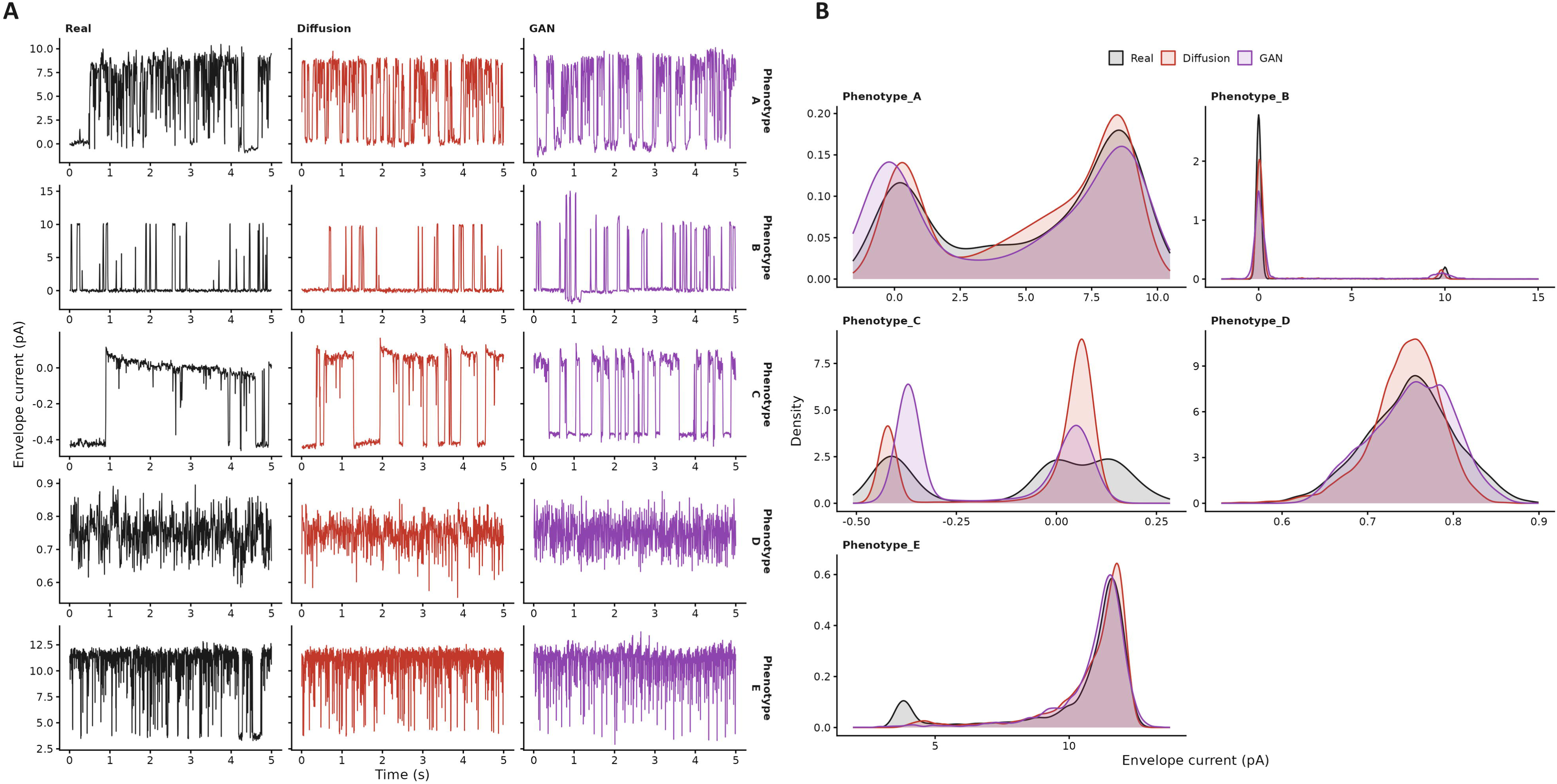
The slow phase. **(A)** Block-averaged envelope at “decimation” factor **f** = 64. Each of the three arms is shown for of each phenotype. **(B)** Amplitude histograms of the decimated coarse envelope/records for the same three arms using kernel density as described above for Figure 6. Real record (black), the diffusion-generated record (red) and the DeepGANnel-generated record (purple).

### 3.6. Amplitude distribution equivalence testing

To complement our qualitative amplitude-histogram comparison (Figure 6), we tested statistical equivalence of the amplitude distribution between each real seed record and its GAN and Diffusion arms, using a one-sided equivalence test (Wellek, 2010) on the Kolmogorov-Smirnov distance between windows of the current trace: the margin (Δ) is the 95th percentile of the KS distance between windows drawn from the real record’s own first and second halves; equivalence requires the real-vs-arm window-pair distribution’s 95th percentile to fall below Δ. Window length was 1024 samples for all phenotypes.

### 3.7. Data Generation

Figure 9 shows generated records with block boundaries marked. Subjectively. there are no discontinuities in amplitude, noise texture or baseline drift in the generated record; a process that has not been possible for windowed based GAN approaches (Ball et al., 2022). To formally test if this join is visible, we compared the distribution of sample-to-sample differences within half a carried context of each join against those elsewhere in a block (with KS tests), to see if these joined simulations are significantly different than points outside of this continuity accommodation. Joins are not detectable in 3 of 5 phenotypes (B p = 0.513, C p = 0.292, E p = 0.095) and marginally detectable in 2 (A p = 0.013, D p = 0.014); KS statistics 0.0050 to 0.0097. This allows practically unlimited record length without a boundary between generated windows, a process that brings neural network approaches in line with non-deep learning simulation approaches.

**Figure 9.**
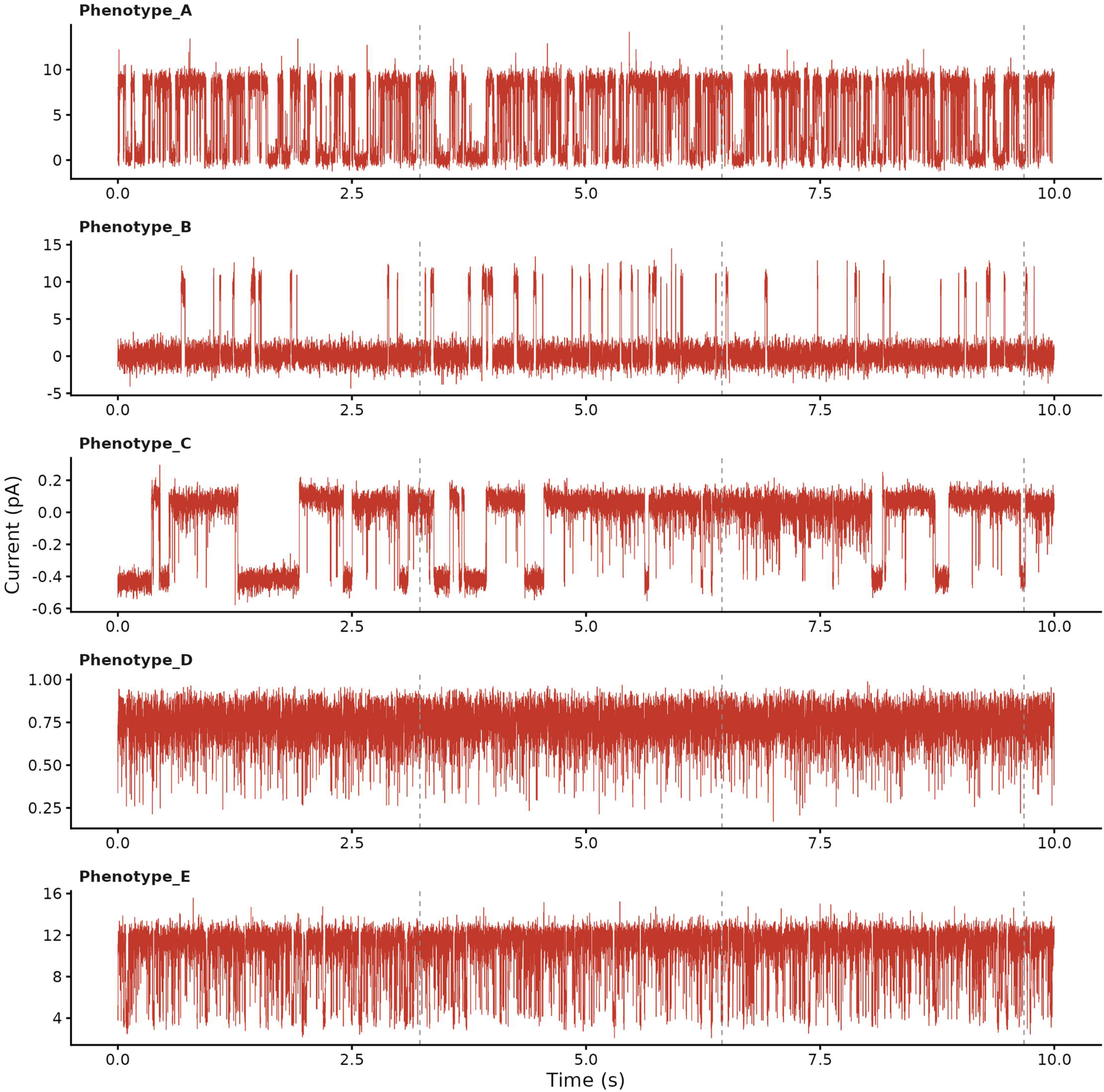
Examples of “Seamless” long-record generation. Since the U-Net is fully convolutional, it has no fixed input length, so it can generate contiguous records far longer than the 128-sample windows used in training, but longer runs become computationally intensive and are limited by available GPU memory. Therefore we generate consecutive blocks joined by RePaint context-carry inpainting (see Methods). We show the first 10 s (100,000 samples) of the generated record here for each phenotype (decimated for display). Faint vertical dashed lines mark the three block boundaries within this stretch. Blocks are 32,768 samples long with a carried context of 512 samples. Each full 200 s (2,000,000-sample) record contains 61 joins, all of which enter the block-join test.

### 3.8. Kinetic analysis

To test for preservation of the kinetic structure of the generated records, we plot the dwell-time distributions for generated and seed records in Figure 10. Burst structure was compared in the same way (Figure 11), using a critical shut time (*Tcrit*) fitted to each real seed recording (Colquhoun and Sakmann, 1985) and thence applied to all three arms; phenotypes B and D yielded no critical time and are therefore not shown. As the idealisation was simulated from a specified transition probability matrix, we know the expected dwell-time distribution *a priori*. For the diffusion network, this simulation is a conditioning signal as an input to the model, whereas for GAN approaches this is simply an output.

**Figure 10.**
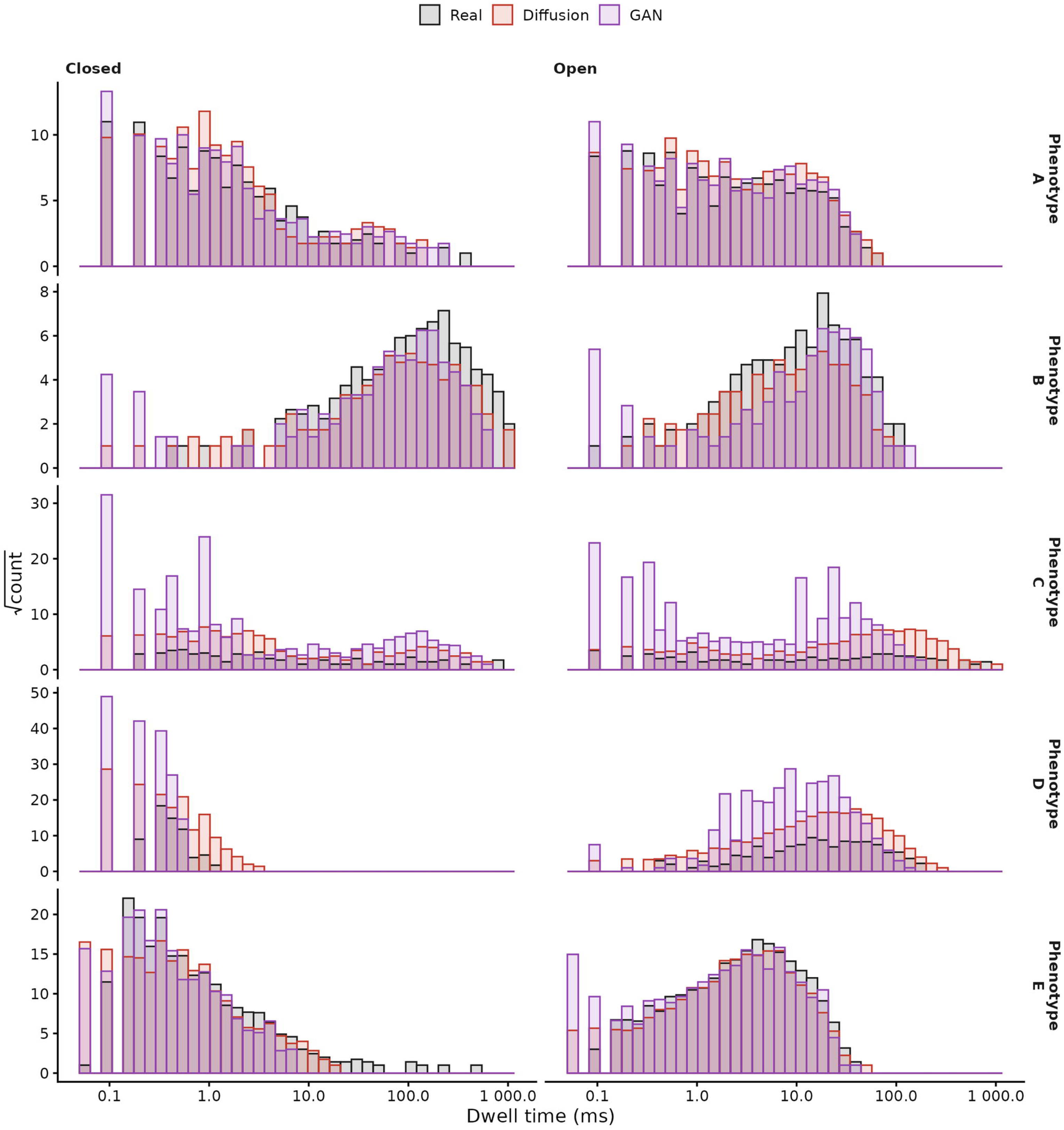
Dwell times from each arm’s Channels column. Closed and open dwell-time distributions taken from the Channels column of each arm (Real, Diffusion, GAN), log-binned with square-root ordinate (Sigworth and Sine, 1987). For the diffusion arm this column is the exact conditioning ground truth; for DeepGANnel it is a network output. Note this measurement fails (in DiffuChannel) in cases where, the dwell times are not practicably recoverable from the noisy (“raw”) current, for example with the rapid kinetics of Phenotype D.

**Figure 11.**
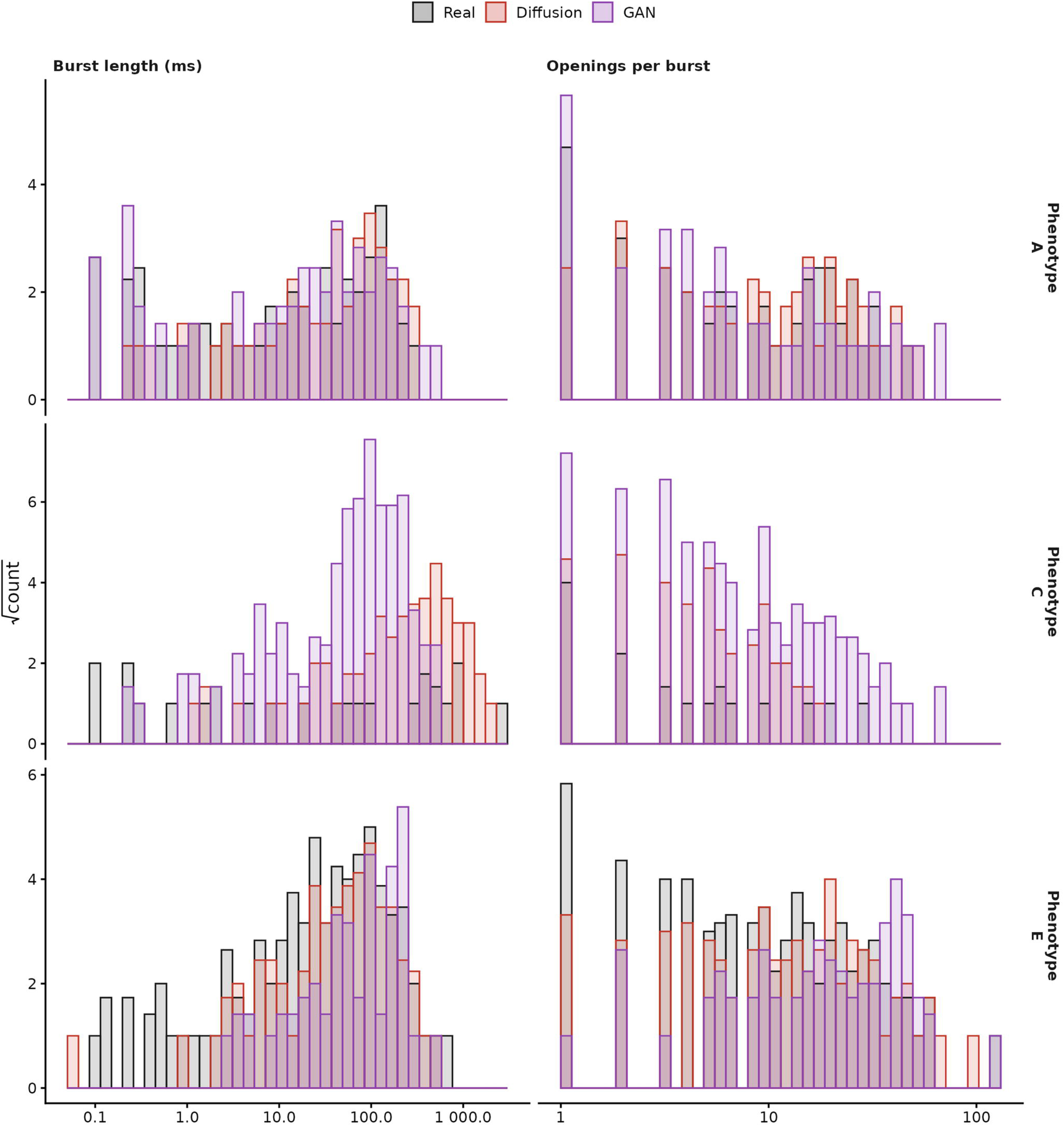
Burst kinetics. Burst kinetics for the three phenotypes; based on an estimated critical time (*Tcrit*) which defines a burst. Where these were derivable from the data (note that Phenotypes B and D are not shown since they yielded no *Tcrit* because their closed times were effectively monoexponential). The left column shows burst lengths in milliseconds and the right column shows number of openings per burst. Both have log abscissa with square-root ordinate, as per Sigworth and Sine (Sigworth and Sine, 1987). *Tcrit* was fitted to each phenotype’s real record by the Colquhoun-Sakmann “missclassified events” criterion (Colquhoun and Sakmann, 1985) and then imposed across all three arms (for consistency).

#### 3.8.1. Gating-kinetics equivalence testing

We further tested statistical equivalence of the noise level, open probability, and dwell-time distributions between each real record and its GAN and Diffusion arms, all as one-sided equivalence tests (Wellek, 2010) with the same real-split-half margin construction as above. Noise SD was measured per qualifying open/closed event (≥30 consecutive samples in one state) rather than fitted, giving a natural distribution of noise levels with no fitting assumption. Open probability was measured per window (1024 samples) rather than as a single global figure (Table 6). Dwell-time equivalence was tested on the full duration distribution per state (Open, Closed), not merely the mean dwell time (Tables 7). Open probability was found equivalent for the diffusion arms in two of the five phenotypes (A and C) but only one for DeepGANnel (A), and dwell-time distributions in four of ten phenotype/state combinations against three, respectively (Tables 6 and 7).

**Table 6.**
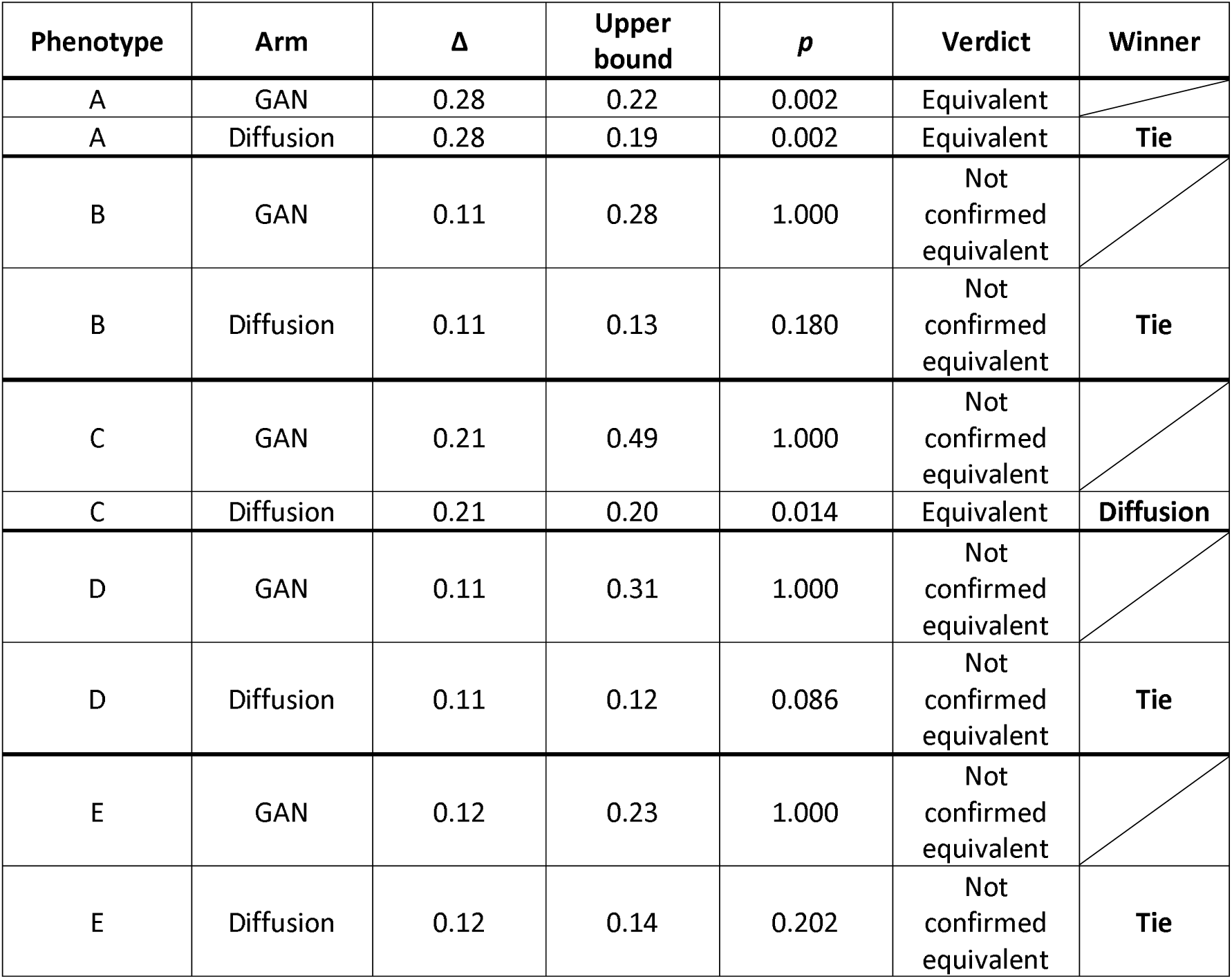
Open-probability equivalence. Same one-sided procedure, Δ/*Upper-bound*/*Verdict* logic and Winner criterion as Table 4, applied to open probability measured *per 1024-sample window*.

**Table 7.** Dwell-time distribution equivalence. Same one-sided procedure as Tables 3, 4 and 6, applied to the full open/closed dwell-time duration distribution (individual event durations, *ms*). *n* (real/arm) the total number of dwell events contributing to real and to the respective arm.

| Phenotype | Arm | State | n<br>(real/arm) | $\Delta$ | Upper<br>bound | $p$ | Verdict | Winner |
| --- | --- | --- | --- | --- | --- | --- | --- | --- |
| A | GAN | Open | 899/1030 | 0.15 | 0.08 | 0.002 | Equivalent |  |
| A | Diffusion | Open | 899/1100 | 0.15 | 0.11 | 0.004 | Equivalent | Tie |
| A | GAN | Closed | 900/1029 | 0.10 | 0.09 | 0.006 | Equivalent |  |
| A | Diffusion | Closed | 900/1100 | 0.10 | 0.13 | 0.365 | Not confirmed equivalent | GAN |
| B | GAN | Open | 466/335 | 0.14 | 0.22 | 0.850 | Not confirmed equivalent |  |
| B | Diffusion | Open | 466/282 | 0.14 | 0.18 | 0.261 | Not confirmed equivalent | Tie |
| B | GAN | Closed | 467/335 | 0.17 | 0.23 | 0.719 | Not confirmed equivalent |  |
| B | Diffusion | Closed | 467/283 | 0.17 | 0.20 | 0.224 | Not confirmed equivalent | Tie |
| C | GAN | Open | 139/2850 | 0.48 | 0.37 | 0.002 | Equivalent |  |
| C | Diffusion | Open | 139/681 | 0.48 | 0.33 | 0.002 | Equivalent | Tie |
| C | GAN | Closed | 140/2850 | 0.29 | 0.45 | 1.000 | Not confirmed equivalent |  |
| C | Diffusion | Closed | 140/681 | 0.29 | 0.18 | 0.002 | Equivalent | Diffusion |
| D | GAN | Open | 818/6657 | 0.13 | 0.30 | 1.000 | Not confirmed equivalent |  |
| D | Diffusion | Open | 818/3170 | 0.13 | 0.08 | 0.002 | Equivalent | Diffusion |
| D | GAN | Closed | 817/6656 | 0.12 | 0.55 | 1.000 | Not confirmed equivalent |  |
| D | Diffusion | Closed | 817/3170 | 0.12 | 0.37 | 1.000 | Not confirmed equivalent | Tie |
| E | GAN | Open | 2894/2967 | 0.06 | 0.15 | 1.000 | Not confirmed equivalent |  |
| E | Diffusion | Open | 2894/2584 | 0.06 | 0.07 | 0.110 | Not confirmed equivalent | Tie |
| E | GAN | Closed | 2895/2966 | 0.07 | 0.12 | 1.000 | Not confirmed equivalent |  |
| E | Diffusion | Closed | 2895/2584 | 0.07 | 0.17 | 1.000 | Not confirmed equivalent | Tie |

### 3.9. Manifold projections

Finally, Figure 12 shows t-SNE and UMAP projections of 512 sample windows drawn from diffusion generated and real seed data records, embedded on the same axes. We find that the data is clustered closely with similar pattens for each phenotype. Cluster analysis using ClusterSignificance permutation tests (100 iterations per comparison) found the real-vs-diffusion and real-vs-GAN point clouds were significantly separable (p ≤ 0.05) in 6 of 10 and 4 of 10 phenotype/projection combinations respectively, with a median *p*-*value* of 0.015 across all 30 comparisons (see Table 8).

**Figure 12.**
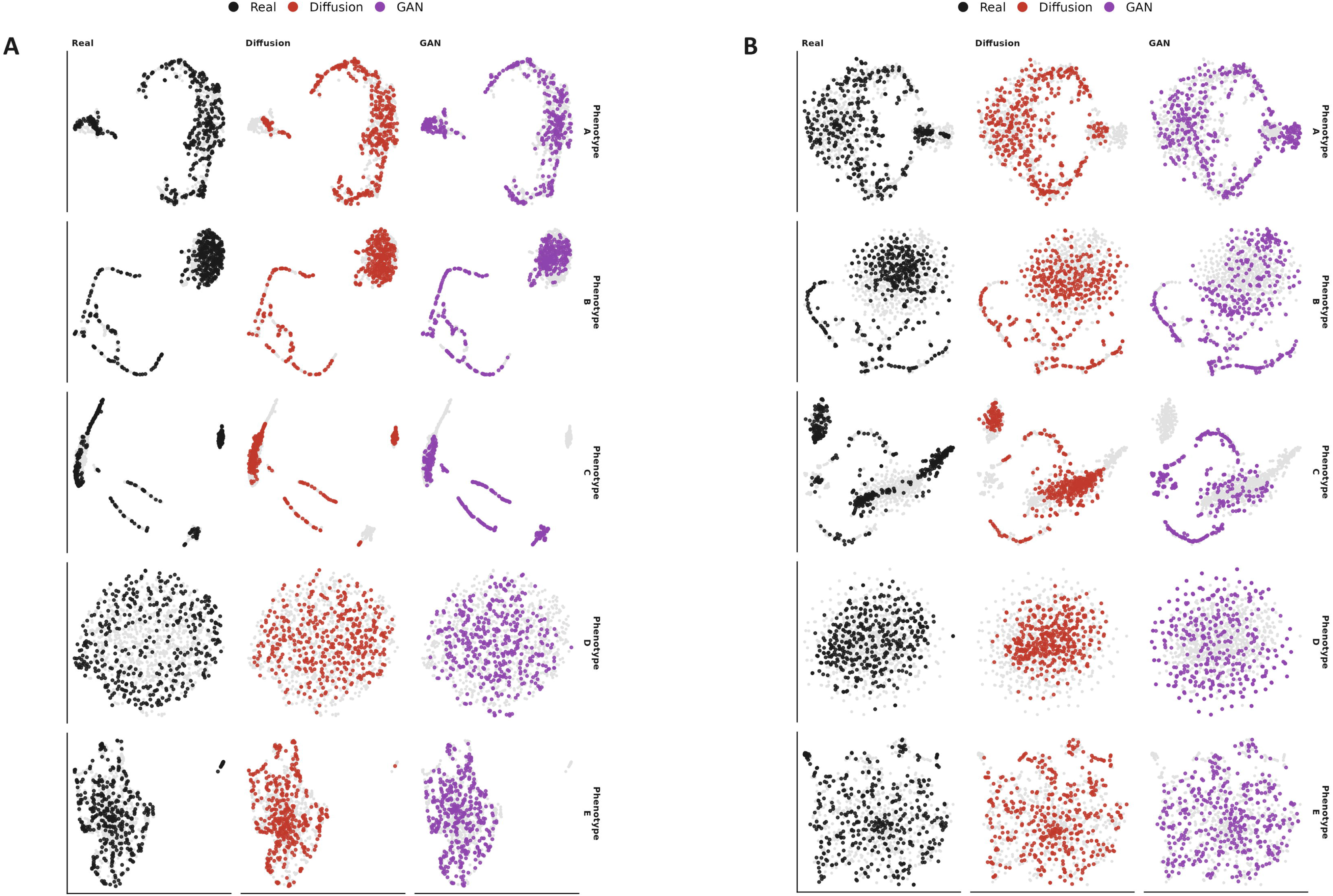
Manifold projections. UMAP **(A)** and t-SNE **(B)** embeddings of 512-point noisy (raw) current windows from all three arms (Real, Diffusion and GAN), embedded jointly within each phenotype and rescaled so the three columns share common axes. Each panel colours one of the arms over the grey “ghost” of the full embedding superset. Quantification of clustering similarity itself is included in Table 8.

**Table 8.** Manifold cluster separation. ClusterSignificance permutation test (Serviss et al., 2017) of whether each synthetic arm’s point cloud can be separated from that of the real record in the UMAP and t-SNE embeddings of Figure 12, run separately for each phenotype and projection with 100 permutations. *p* is the fraction of permutations giving a separation score at least as large as the true labels; with 100 permutations the smallest reportable value is 0.01, shown as ≤0.01. Unlike Tables 3, 4, 6 and 7, a large *p* is the favourable result: it means the synthetic and real point clouds. *p-*values of larger than 0.05 indicate that the synthetic and real are not statistically significantly different. This does not mean “the same”, and we awarded winner to the arm with the largest value.

| Phenotype | Projection | Arm | *p* | Winner* |
| --- | --- | --- | --- | --- |
| A | UMAP | GAN | 0.31 |  |
| A | UMAP | Diffusion | 0.11 | GAN |
| A | t-SNE | GAN | 0.12 |  |
| A | t-SNE | Diffusion | 0.24 | Diffusion |
| B | UMAP | GAN | $\leq 0.01$ | |
| B | UMAP | Diffusion | 0.03 | Diffusion |
| B | t-SNE | GAN | $\leq 0.01$ | |
| B | t-SNE | Diffusion | $\leq 0.01$ | Tie |
| C | UMAP | GAN | $\leq 0.01$ | |
| C | UMAP | Diffusion | $\leq 0.01$ | Tie |
| C | t-SNE | GAN | $\leq 0.01$ | |
| C | t-SNE | Diffusion | $\leq 0.01$ | Tie |
| D | UMAP | GAN | 0.46 |  |
| D | UMAP | Diffusion | $\leq 0.01$ | GAN |
| D | t-SNE | GAN | 0.37 |  |
| D | t-SNE | Diffusion | 0.30 | GAN |
| E | UMAP | GAN | 0.74 |  |
| E | UMAP | Diffusion | $\leq 0.01$ | GAN |
| E | t-SNE | GAN | 0.37 |  |
| E | t-SNE | Diffusion | 1.00 | Diffusion |

\*In this case “winner” denotes the higher *p-value*.

Finally, Figure 13 summarises the similarity metrics of Table 5 in terms of within-phenotype pairs. The diffusion arm scored better than DeepGANnel in 42 of the 55 phenotype–metric comparisons, and in every phenotype for ACF RMS, mean closed dwell time and open probability.

**Figure 13.**
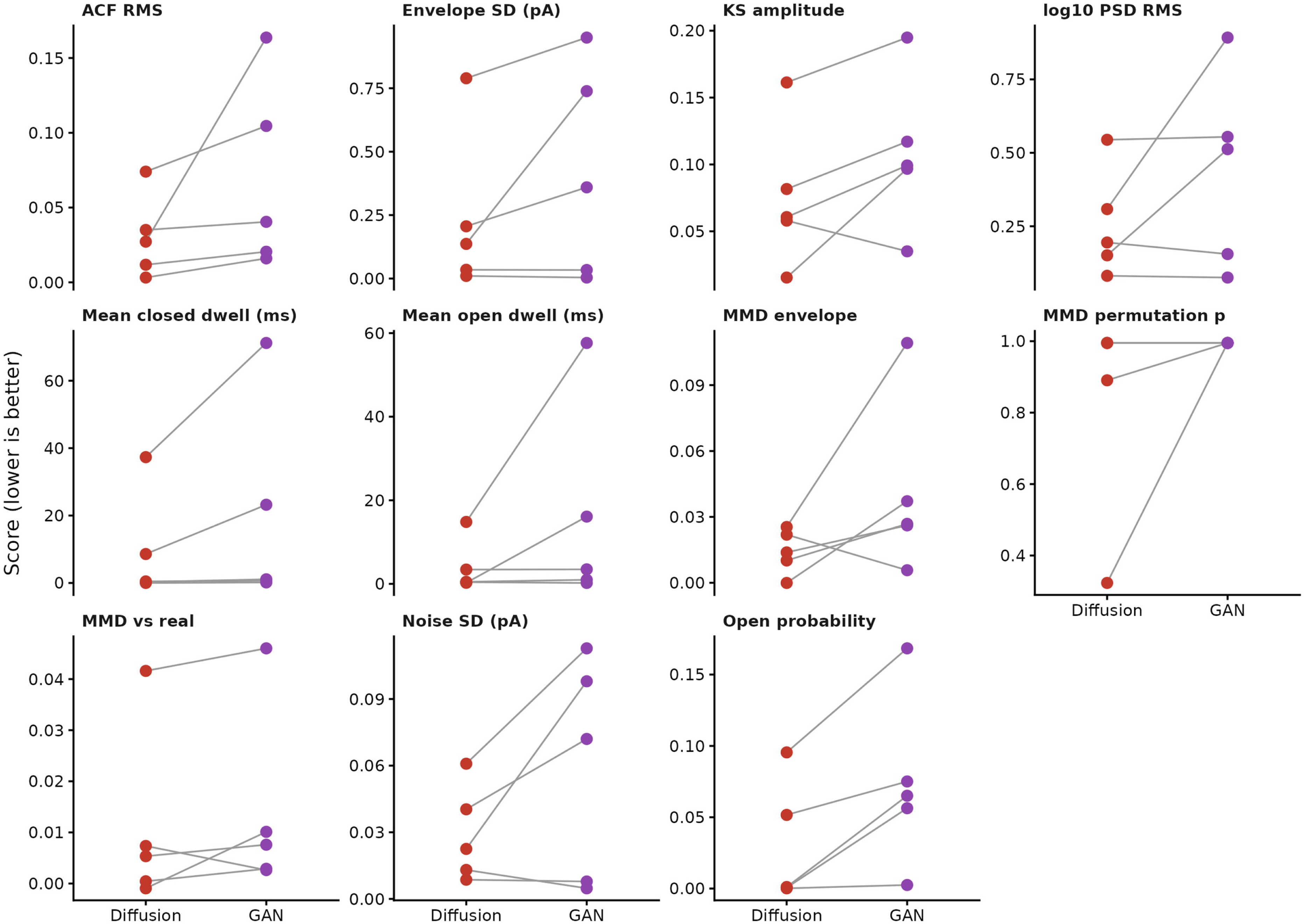
Paired metric summary. A summary of similarity-metrics, as detailed in Table 5, each shown as within-phenotype pairs, diffusion against DeepGANnel; each grey line joins the two arms’ scores for one phenotype. Scores are oriented so that lower is better throughout: the MMD permutation p is plotted as 1 − p, and metrics that should match the real record (noise SD, envelope SD, open probability, mean dwell times) as the absolute difference from the real value.

## 4. Discussion

Traditional single ion-channel simulation relies on mathematical and signal processing approaches in order to preserve the underlying structure of the data; however, struggles to generate some of the artefacts and noise present in real lab-sourced recordings. Generative AI approaches (e.g. DeepGANnel) can generate convincing artefacts and noise, but may contain errors with the underlying mathematical structure and suffer from a lack of continuity between generated windows. In this work, we have demonstrated that diffusion models allow for a combination of the best parts of both of these approaches: continuous, robust and well-defined underlying simulation from predefined transition probability matrices, as well as convincing noise and artefacts we would see in lab-recorded data.

### 4.1. Robust Paired Ground Truth

For downstream application of simulated data, the association with an accurate paired ground truth is essential. Traditional methods excel in this area at a cost of realism, whereas generative AI approaches (as highlighted by Oikonomou et al (Oikonomou et al., 2024)) obscure the source of this ground truth; and it is not known whether these labels are accurate enough to train a downstream model.

In this work, the idealisation acts as a conditioning *input* to the model rather than a predicted output. Therefore, the output idealisation is always generated mathematically and holds the structure of the underlying Markovian structure. We believe that this will allow more robust downstream model development under testing of a wider variety of synthetic channel phenotypes.

### 4.2. Comparison to Adversarial Approaches

Diffusion models, compared to adversarial GAN based approaches, do not exhibit the phenomenon of “modal collapse”, and are generally much simpler to train. In training DeepGANnel (Ball et al., 2022); manual learning rate adjustment of the models was required throughout training to ensure a good balance between the discriminator and generator models remained. I f one model performed at a far greater accuracy than the other, we found that training would fail to recover, even if learning rates were adjusted. For our diffusion model presented here, no training supervision is required, as the model is simply minimising the mean squared error of the outputs with no adversarial loss. This results in faster end-to-end training, and greater stability.

Generation time is longer for our diffusion model than the previous GAN. This is because GAN model data generation rely only on the generator model with a single forward pass to produce a new data record. DDPM models require several sequential network passes of a larger model to produce a single record. We find that to produce 10 seconds of data (post training), the diffusion model took on average 6.1 seconds to generate a record, whereas DeepGANnel took roughly 4ms per record. Various approaches exist can speed up the diffusion generation, but we did not test these (Song et al., 2021; Rombach et al., 2022; Salimans and Ho, 2022).

### 4.3. Long-Term Feature Recovery

Perhaps the most important contribution of this work is the generation of realistic long-term noise terms such as baseline drift, seal degradation or bursting in the generated record. While mathematical approaches exist to approach modelling these (e.g. low order ^n^ noise), they rarely result in accurate simulations of these phenomena. Our approach of supplying a coarse signal as additional conditioning to the record is a relatively simple way of approaching this (Dhariwal and Nichol, 2021); and the same approach could apply to ECG, EEG, nanopore or other electrophysiological data; all of which have long-timescale structures present in long recordings. The “dwell” and “burst” clocks as part of this coarse envelope simulation also address this (Vaswani et al., 2017). Through inclusion of this information into the conditioning signal, the diffusion model can model within-event artefacts (e.g. relaxation) that would otherwise be impossible for events longer than a window size. This conditioning information is all derived from the simulated idealisation, and therefore does not constitute a data leakage risk for our evaluation methods.

### 4.4. Limitations

We have benchmarked against our existing DeepGANnel work, but not against mathematical models. We avoid benchmarking against Gillespie (Gillespie, 1977) style simulations due to sensitivity of simulation parameters to simulation performance. Poor simulation performance for these methods can simply be due to poor handcrafting, for example, the user did not “try hard enough” to make it perfect, or just because the method really is inadequate. Therefore, we omitted inclusion of Gillespie from our analyses here.

While DiffuChannel shows greater performance cross several metrics compared to DeepGANnel, it often does not do so significantly, and on some occasions, it is beaten by GAN. Nevertheless, we believe that many aspects of DiffuChannel’s design (e.g. ease of use, continuous generation with ensured continuity and most importantly, training stability) justify the importance of this work. As a point of note we ourselves will definitely switch to this method of simulation/augmentation from DeepGANnel now.

We also acknowledge that training and evaluation is limited to a small number of channel phenotypes and data. A lack of training data within a channel phenotype limits the availability of features for the model to produce; a model cannot reproduce any recording artefacts that are not part of the training data. DeepGANel also exhibited this problem (Ball et al., 2022), however we are keen to evaluate our method across a larger range of channel phenotypes and behaviour in the future to test our method’s robustness.

The methods we use here process data at time points, therefore, it should be noted that any events (or features) shorter than the sample rate itself will be lost from the idealisation. This is actually common in the digital age of electrophysiology where even raw data are typically sampled at discrete time points, even if this is at a very high sample rate. In fact this limitation also exists in analysis tools such as our own DeepChannel (Celik et al., 2020) and even QuB (Nicolai and Sachs, 2013). These would be unable to recover such transitions in data from any source in any case.

### 4.5. Future directions

We invite developers to test DiffuChannel and comment on how effective they find this model – and to use these data in development of further single-channel analysis methods. Subject to available time, we may also be able run this for users on request sometimes.

## 5. Data and code availability

Code: https://github.com/RichardBJ/DiffuChannel. Seed and generated datasets (modest to meet GitHub’s fair use principles). The analysis notebook reproducing every figure and metric is included in the repository.

## 8. Ethics

No new animal or human data were used in this study.

